# Discordant results among MHC binding affinity prediction tools

**DOI:** 10.1101/2022.12.04.518984

**Authors:** Austin Nguyen, Abhinav Nellore, Reid F. Thompson

## Abstract

A large number of machine learning-based Major Histocompatibility Complex (MHC) binding affinity (BA) prediction tools have been developed and are widely used for both investigational and therapeutic applications, so it is important to explore differences in tool outputs. We examined predictions of four popular tools (netMHCpan, HLAthena, MHCflurry, and MHCnuggets) across a range of possible peptide sources (human, viral, and randomly generated) and MHC class I alleles. We uncovered inconsistencies in predictions of BA, allele promiscuity and the relationship between physical properties of peptides by source and BA predictions, as well as quality of training data. Our work raises fundamental questions about the fidelity of peptide-MHC binding prediction tools and their real-world implications.

## INTRODUCTION

Human Leukocyte Antigen (HLA) alleles are critical components of the immune system’s ability to recognize and eliminate tumors and infections (1). Infectious diseases in particular are thought to be a major source of selective pressure on the Major Histocompatibility Complex (MHC) region which encodes HLA alleles and is one of the most diverse regions of the human genome (2–8). There is large diversity in the antigenic peptide sequences which individual HLA alleles can recognize and ultimately present to the adaptive immune system (9), with a positive correlation between increased sequence diversity recognition and fitness (10).

Tools that can predict the extent to which a given HLA allele may have an affinity for a given peptide have critical implications for our ability to understand and translationally leverage antigen-specific immune response pathways. For instance, MHC binding affinity predictors have been – or otherwise have the potential to be – used to evaluate an individual or population’s susceptibility to viral infection (11), to develop an understanding of specific autoimmune conditions (12), to improve transplantation technologies (13), or even to assist in the development of personalized cancer vaccines (14–18). Numerous peptide-MHC binding prediction tools exist, and are key components in broader antigen prediction methodologies (19–22).

The most widely adopted MHC binding prediction tools rely on neural network models trained on binding affinity (BA) and/or eluted ligand (EL) data. The most commonly cited tool, netMHCpan (23,24), uses both BA and EL data in a neural network architecture with a single hidden layer to predict allele-specific binding affinities. MHCflurry (25) attempts to improve upon netMHCpan by increasing the number of hidden layers and augmenting BA and EL training data with unobserved decoys. MHCnuggets (26) again trains on BA and EL data but uses a different architecture, with a long short-term memory layer and a fully connected layer to improve its predictions further across different peptide lengths. Lastly, HLAthena (27), while most similar in architecture to netMHCpan, relies on independently generated EL data from mono-allelic cell lines for training.

We sought to better characterize the outputs of these tools over a large and diverse set of peptides, across different tools and HLA alleles, as well as quantify the stability of these predictions. We also sought to measure allelic binding preferences and whether they may enrich for foreign v. self peptides. In this study, we performed a comprehensive *in silico* analysis of peptides from multiple viral proteomes, the human proteome, and randomly generated peptides across HLA class I alleles.

## RESULTS

### Peptide predictions are inconsistent across tools

We first assessed the consistency of peptide-specific MHC I binding affinity predictions across four tools (MHCnuggets, MHCflurry, HLAthena, netMHCpan) and 52 different HLA alleles. We found substantial disagreement in peptide-specific predictions between each tool, independent of allele (Figure 1A), with median intraclass correlation coefficient (ICC) of 0.207 and only 0.48% of peptides having ICC > 0.75. On a per-allele basis, we found a wide range in consistency of predictions across tools, with a mean intraclass correlation as low as 0.12 for A02:07 and as high as 0.64 for A23:01 (Figure 1B). Among all of the peptides predicted by at least one tool to bind to at least one allele, only 7.9% were consistently predicted across all tools to bind to the same allele (Figure 1C).

**Figure 1.**
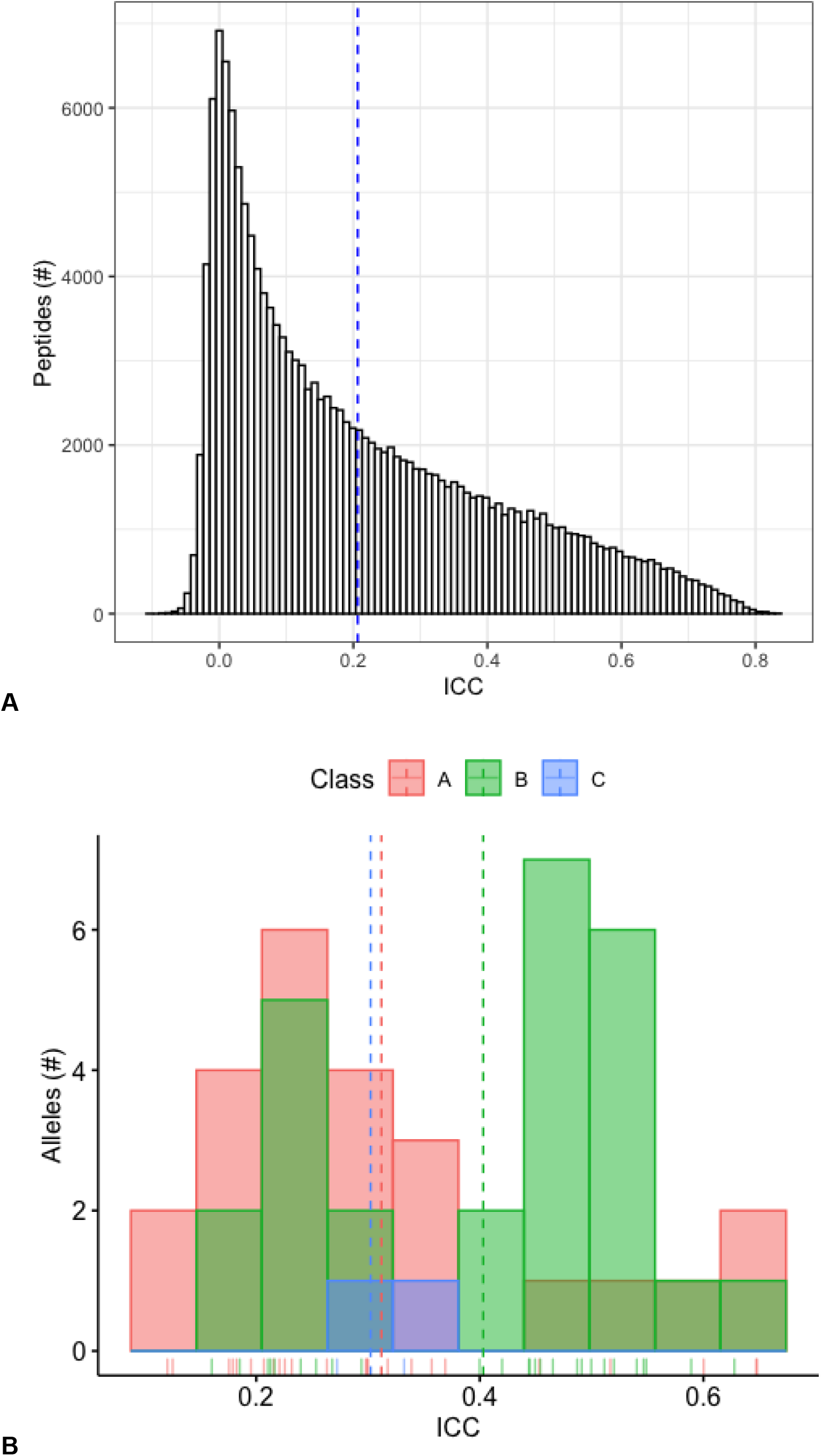
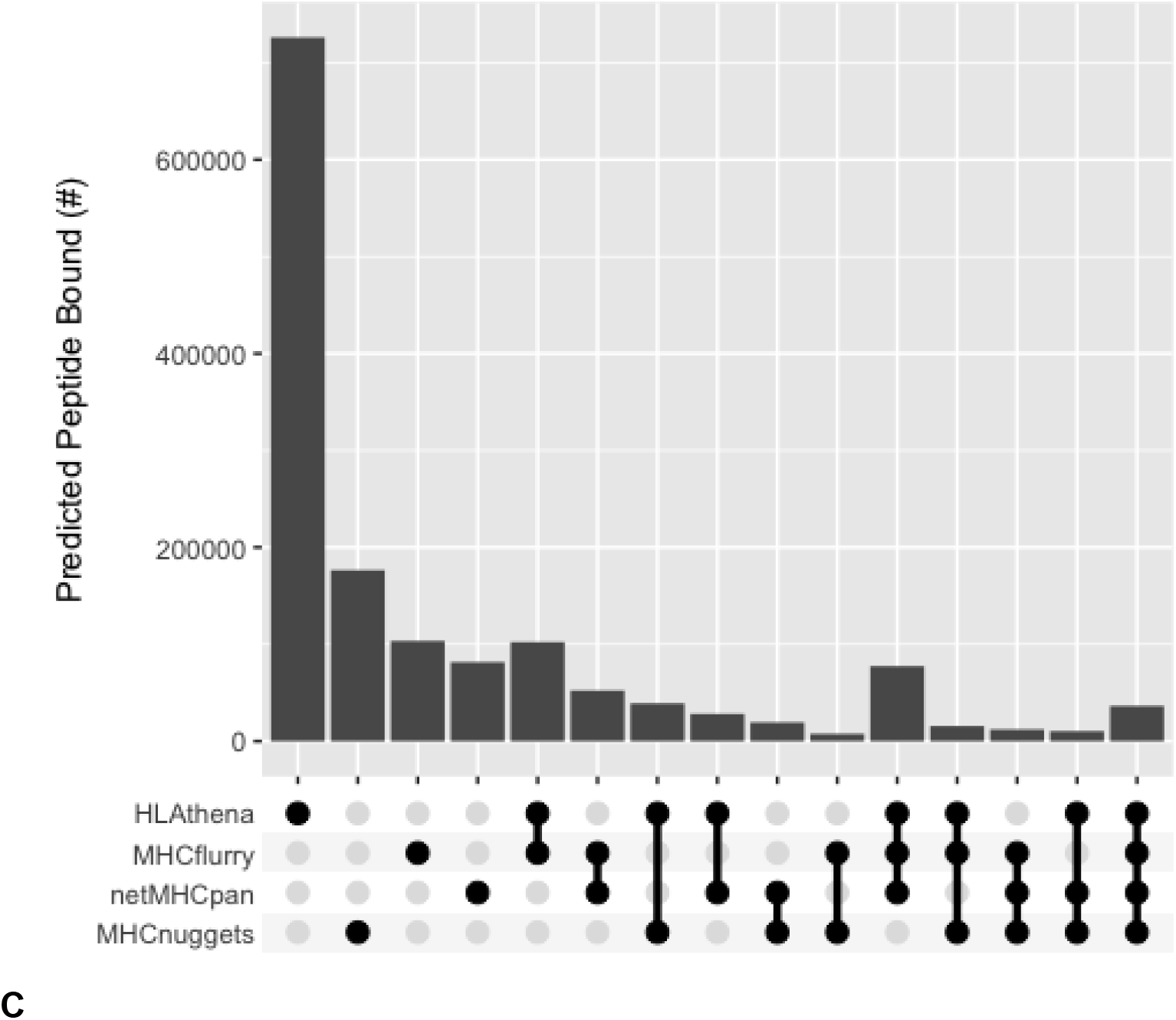
Inconsistency of peptide predictions across tools. A) Histogram of intraclass correlation coefficients (ICC) calculated for a set of 1 million random peptides across four tools (MHCnuggets, MHCflurry, HLAthena, netMHCpan), with ICC calculated as the overall correlation among tools across 52 HLA alleles. The dotted vertical line indicates the median ICC value (0.207) across all peptides. B) Histogram of ICCs for 52 HLA alleles between four tools (MHCnuggets, MHCflurry, HLAthena, netMHCpan). The number of alleles is shown on the y-axis and the ICC is shown on the x-axis. The dotted lines show the mean ICC for alleles belonging to each HLA class. Red, green, and blue colors represent data from -A, -B, and -C alleles, respectively. C) Detailed comparison of the complete set of random peptides predicted to bind (binding score >=0.5) to HLA alleles according to each of four tools. Patterns of agreement or disagreement among groups of peptides predicted by different combinations of tools across 1 million random peptides are shown along each column (e.g. the first column corresponds to peptides predicted by HLAthena while the final column corresponds to peptides predicted by all tools). Each row indicates the predictions associated with the indicated tool. The number of peptides in each column (vertical bars) corresponds to the size of the subset predicted by the indicated combination of tools.

We next investigated aggregate peptide binding predictions across different HLA alleles according to each tool. As others have noted differential HLA allelic promiscuity in peptide presentation (28–31), we too found a wide range in the proportion of peptides a given allele was predicted to bind (Supplementary Figure 1). We uncovered significant inconsistencies in these predictions between tools (Figure 2). Note that this phenomenon is independent of binding affinity threshold (Supplementary Figure 2).

**Figure 2:**
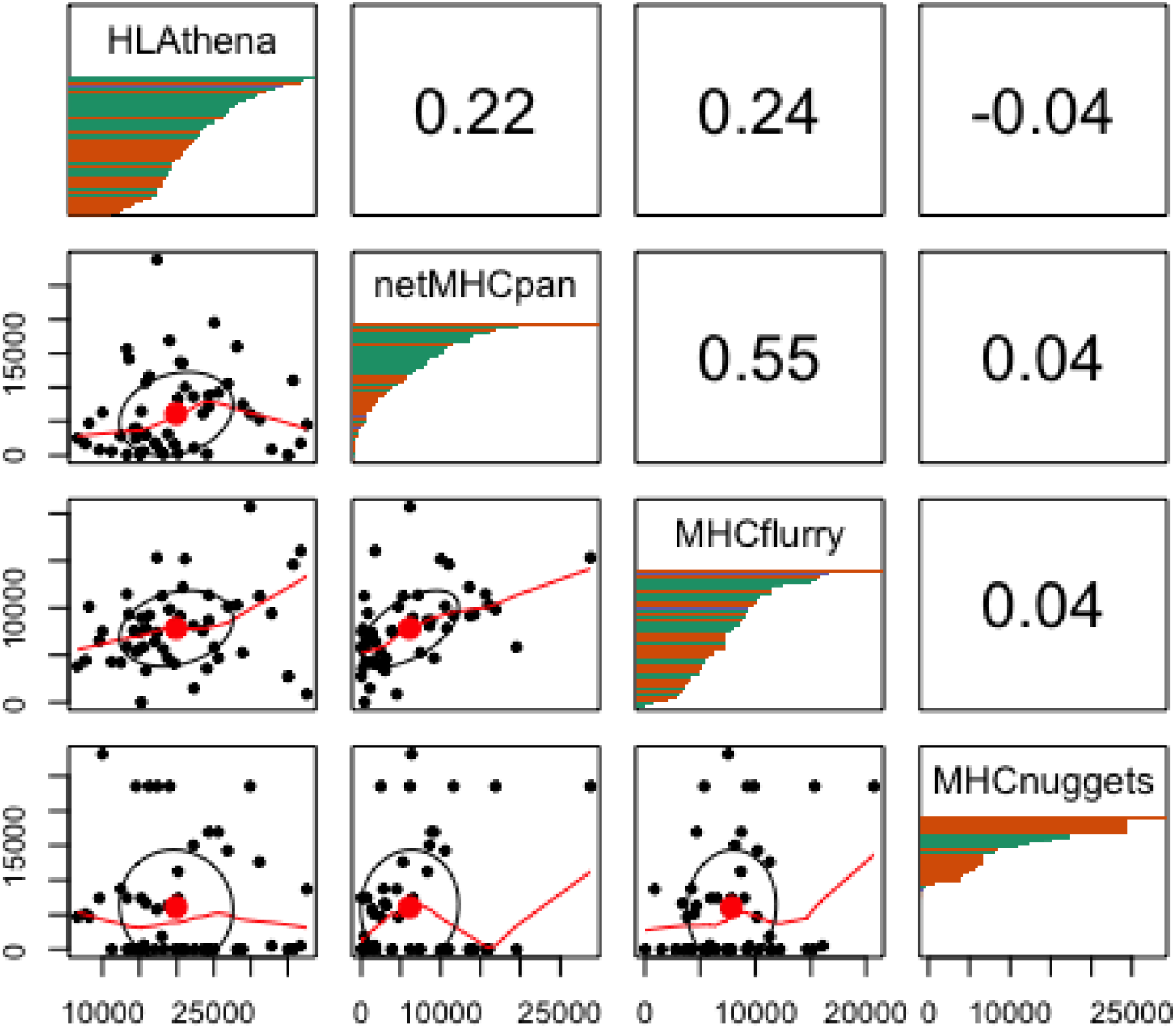
The correlation of HLA allelic presentation of 8-11mers from the random proteome between tools. The lower left grouping of plots displays scatter plots of peptides predicted to bind (>= 0.5 binding probability score) between 2 tools with each point representing the number of predicted binders for each HLA allele. The upper right grouping represents the Spearman correlation of the number of peptides predicted to bind to all alleles between tools. Note that MHCnuggets has a number of alleles with 0 random peptides predicted to bind. The diagonal panels show distribution of HLA allelic presentation from the random proteome for each tool. The number of peptides that putatively bind to each of the HLA alleles is shown along the x-axis as a series of horizontal bars with green, orange, and purple colors representing HLA-A, -B, and -C alleles, respectively, sorted in order of decreasing quantity of binders.

### Amount of training data does not explain inconsistencies between tools

As each allele has a different amount of training data, we were next interested in exploring to what extent the quantity and quality of training data available to each tool might influence its allele-specific predictions. Indeed, some netMHCpan predictive models for some alleles are based on as few as 101 peptides, while others from MHCflurry are based on as many as 31,775 peptides (Supplementary Table 1). Note that we excluded from consideration the ~95% of alleles (4108) that were available for prediction but had no underlying allele-specific training data available (Supplementary Table 2). Ultimately, we found that the amount of training data available was not significantly related to the consistency of binding predictions between tools (Figure 3a), nor was it clearly related to the quantity of binding peptides predicted by tools (Figure 3b).

**Figure 3.**
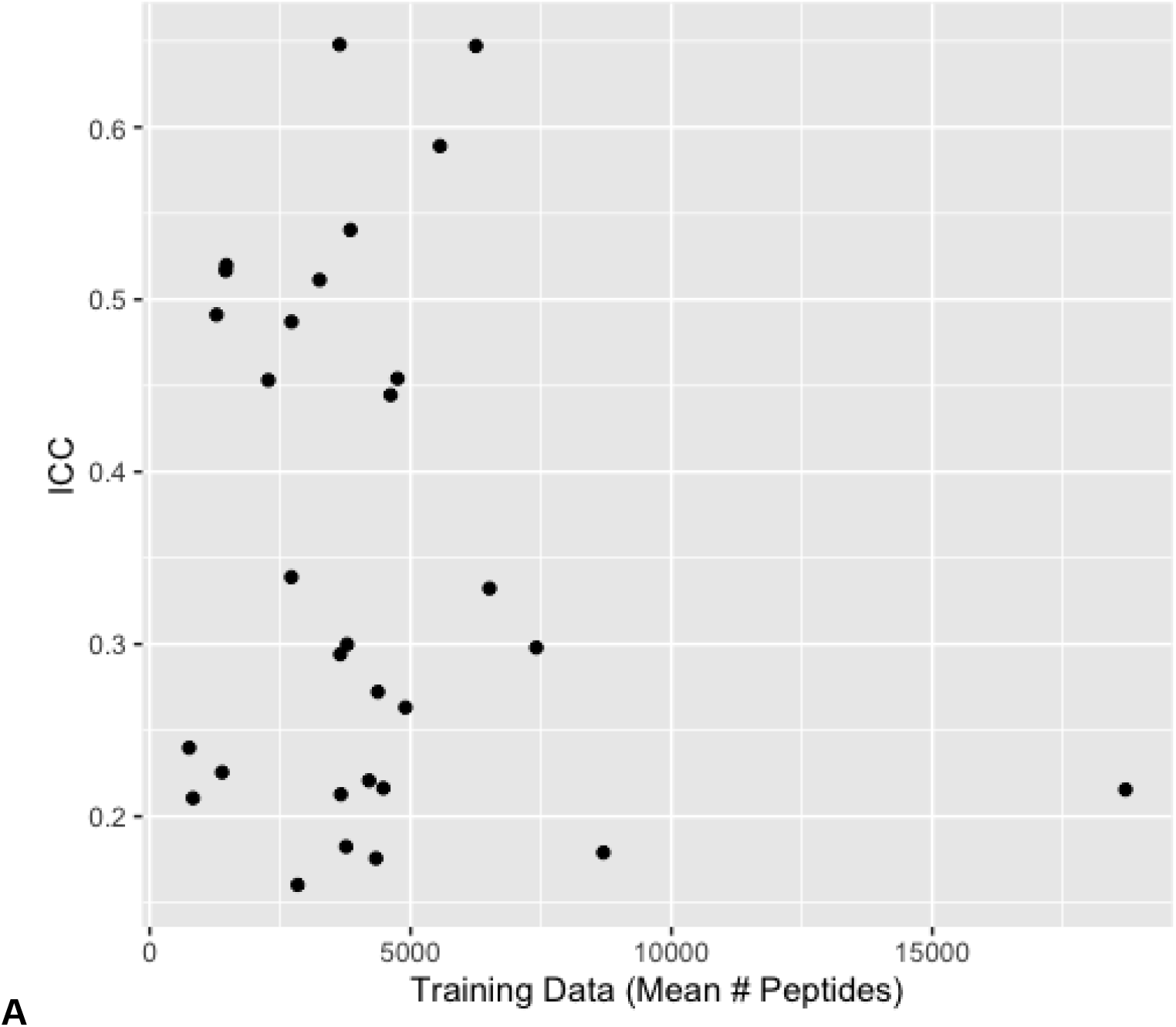
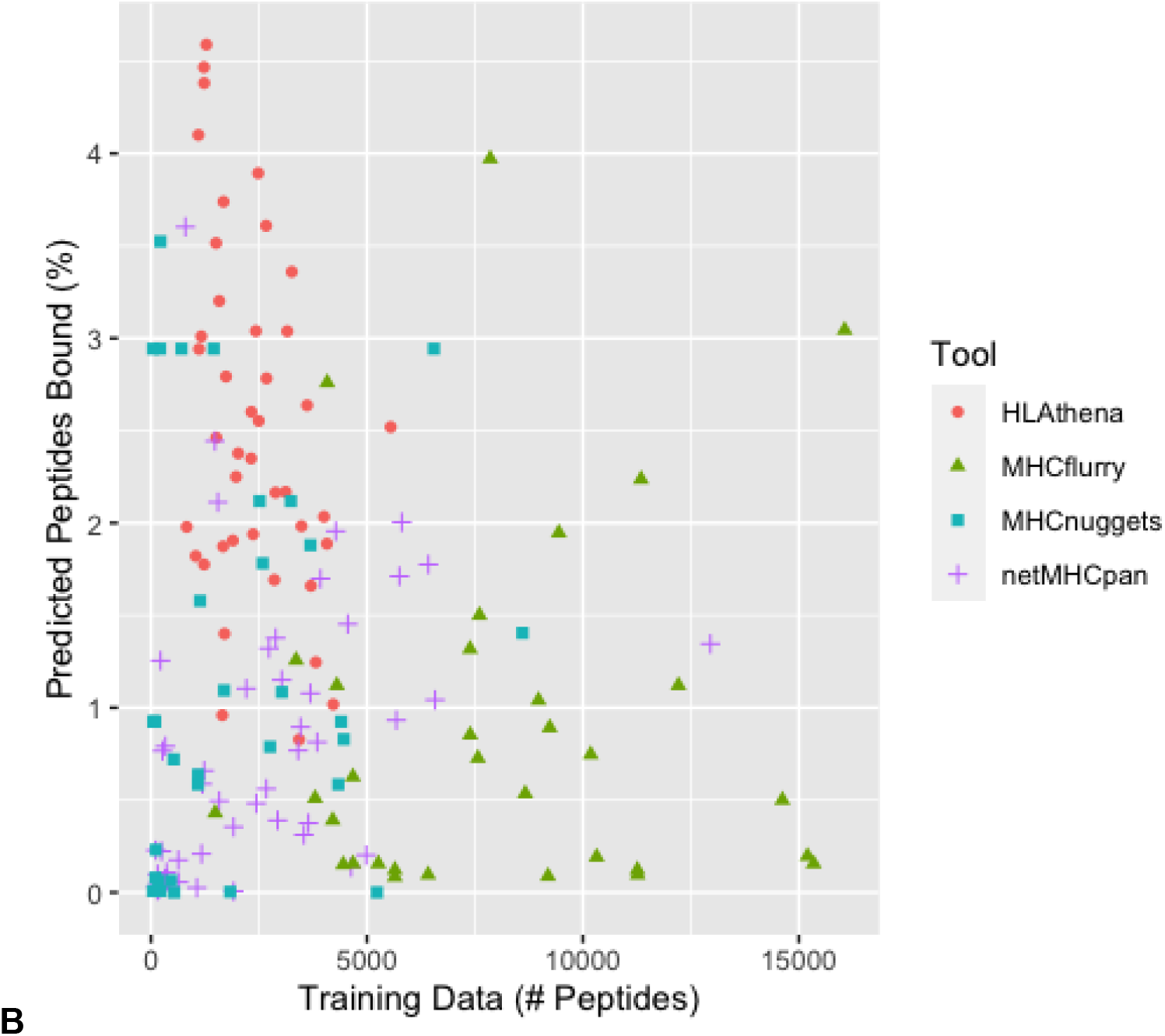
The relationship between training data and consistency of predictions. A) Scatterplot of ICC vs mean training data across 4 tools with each point representing data for a single HLA allele. The mean number of training peptides is shown on the x-axis while the ICC score is shown on the y-axis. B) Scatterplot of the relationship between training data and predicted peptide binding. The number of peptides used as training data for an allele is shown on the x-axis whereas the number of peptides predicted to bind for the same allele is shown on the y-axis. Each dot is a single allele with each color representing a different tool: red circles (HLAthena), green triangles (MHCflurry), blue squares (MHCnuggets), purple plus signs (netMHCpan). We note that netMHCpan does not make all of their training data available, thus the depicted quantity of training data represents an estimate.

### Predicted binding quantities are similar between human and viral proteomes

According to the pathogen driven selection theory of MHC evolution, different HLA alleles are anticipated to be particularly attuned to foreign as opposed to self antigens (3,8,32–35). We therefore sought to compare the predicted capacity of different HLA alleles to present different viral vs. self antigens. Further, we wished to establish which specific alleles had the propensity to bind a larger fraction of peptides in general (allele promiscuity) by observing the relationship between an allele’s ability to bind random peptides versus peptides from a viral or human proteome.

We examined distribution of predicted allelic promiscuity across alleles for 9 sets of peptides of viral, human, and random origin (See Methods). Confining attention to human and viral proteomes, we again found a wide range in the proportion of peptides a given allele was predicted to bind and also significant inconsistencies between tools (Supplementary Figure 3).

We found that the alleles with highest mean binding percentage for human and viral peptides were B15:03 (2.68%) and B15:02 (2.36%) and the allele lowest mean binding percentage were B18:01 (0.24%) and A01:01 (0.33%) (Supplementary Table 3). No alleles were predicted by any tool to preferentially present either viral or human peptides. Further, the distribution of predicted allelic promiscuity across alleles was highly consistent between human and viral proteomes, but not when applied to a set of random peptides (Figure 4). We noted that this phenomenon holds for closely related viruses across all tools and to a lesser extent for more distantly related viruses (Supplementary Figure 4).

**Figure 4.**
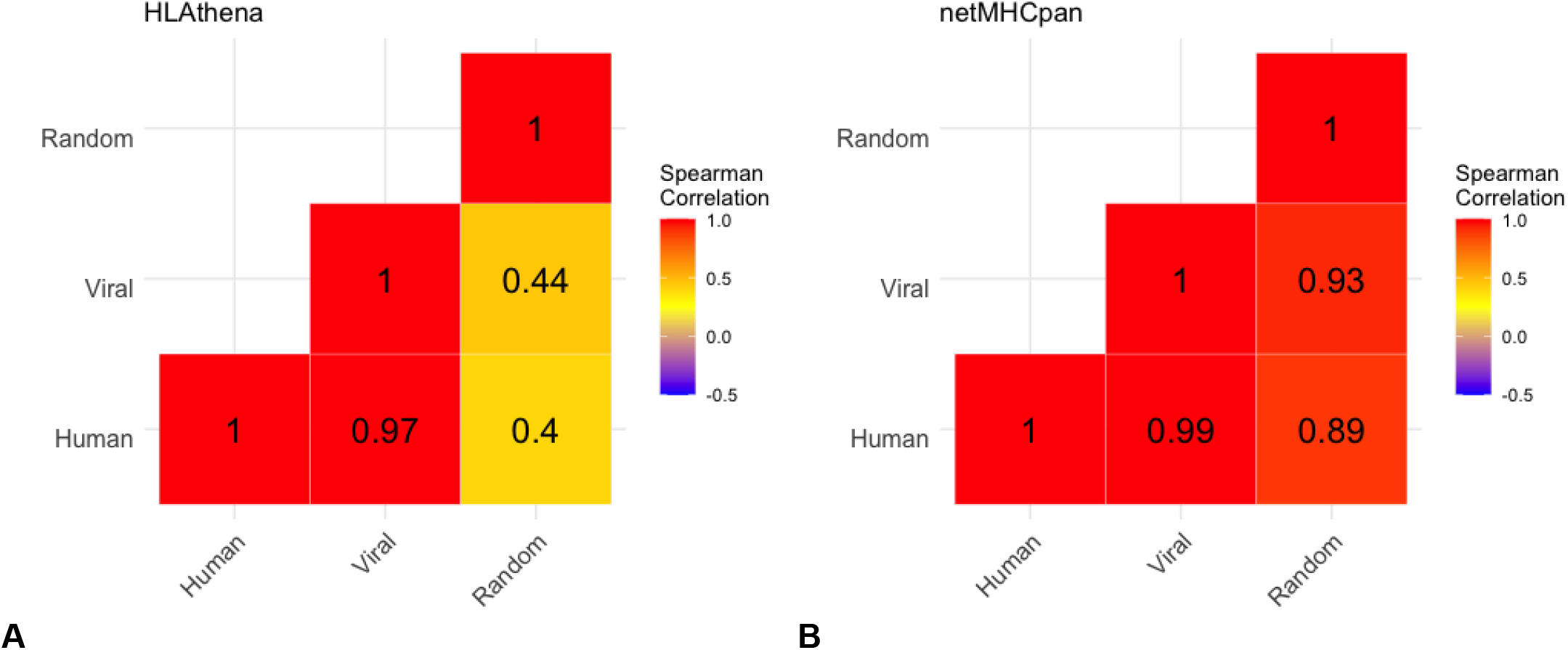
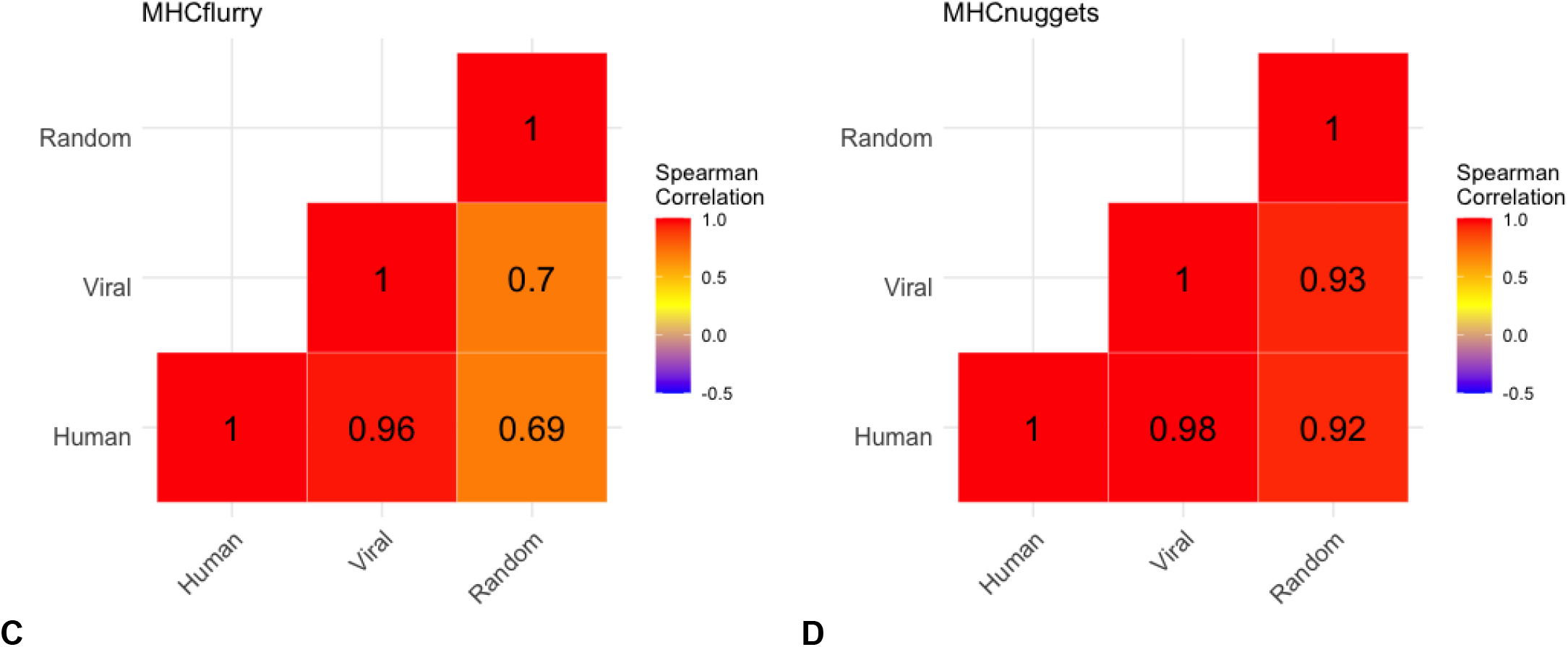
The correlation between peptide sources of predicted allelic promiscuity across alleles. A) Heatmap of spearman correlation between peptide sources for HLAthena-based predictions for human peptides, viral peptides, and randomly generated peptides. Numbers show Spearman correlation coefficients between each pair respectively, while color reflects the Spearman correlation with red approaching a Spearman correlation of 1. Analogous data is shown for netMHCpan, MHCflurry, and MHCnuggets in panels B, C, and D, respectively.

Confining attention to the 9 alleles whose predictive models were likely most robust (based on a minimum of 2000 training peptides for every tool), we again found that the distribution of predicted allelic promiscuity across alleles was consistent between closely related viruses and to a lesser extent between more distantly related viruses (Supplementary Figure 5).

### Peptide physical properties are associated with allele-specific binding predictions

Reasoning that differences in peptide characteristics were the likeliest explanation for predicted differences in binding affinity between different alleles and peptide sources, we next studied the distribution of physical properties among different peptide sets. Human, viral, and random peptide sets all exhibited the same range of physical properties, but were differentially enriched among different physical properties (Supplementary Figure 6). Between individual peptide sets, the differential enrichment ranged from 10% (CMV v. human) to 63% (BK v. random) of peptides (Supplementary Figure 7).

We next sought to discover the relationship between the peptide similarity in physical property space and distribution of predicted allelic promiscuity across alleles. Across all tools, there was a positive relationship between similarity in physical property space and distribution of predicted allelic promiscuity across alleles as evidenced by the negative correlation between peptide set difference and Spearman correlation coefficient (Figure 6).

**Figure 6.**
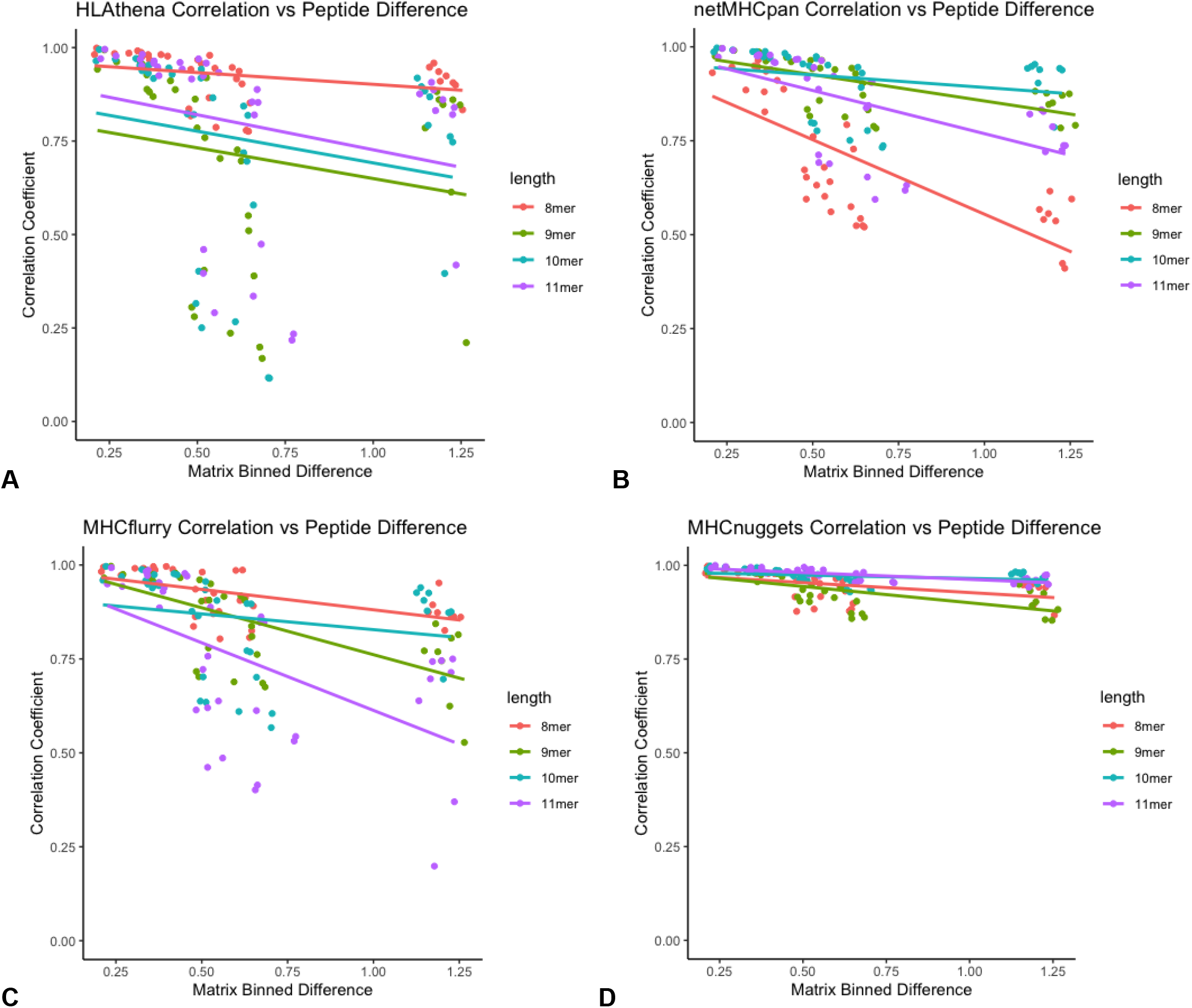
The relationship between physical property similarity vs peptide binding similarity. A) Scatterplot for HLAthena-based predictions, where each point represents predictions for a species vs species pair. Peptide dissimilarity is shown on the x-axis, whereas Spearman correlation coefficients of predicted allelic promiscuity across alleles. Color represents the length of peptide, with 8-, 9-, 10-, and 11-mers shown in red, green, blue and purple, respectively. Analogous data is shown for netMHCpan, MHCflurr, and MHCnuggets in panels B, C, and D, respectively.

Next, we found that each allele has distinct preferences for different peptide physical properties, independent of peptide length (Figure 7A, Supplementary Figure 8). Some alleles (e.g. A01:01 and B08:01) have stronger preference for certain physical properties (Figure 7B,C), while others (B45:01) do not have as clear of a preference (Figure 7D).

**Figure 7.**
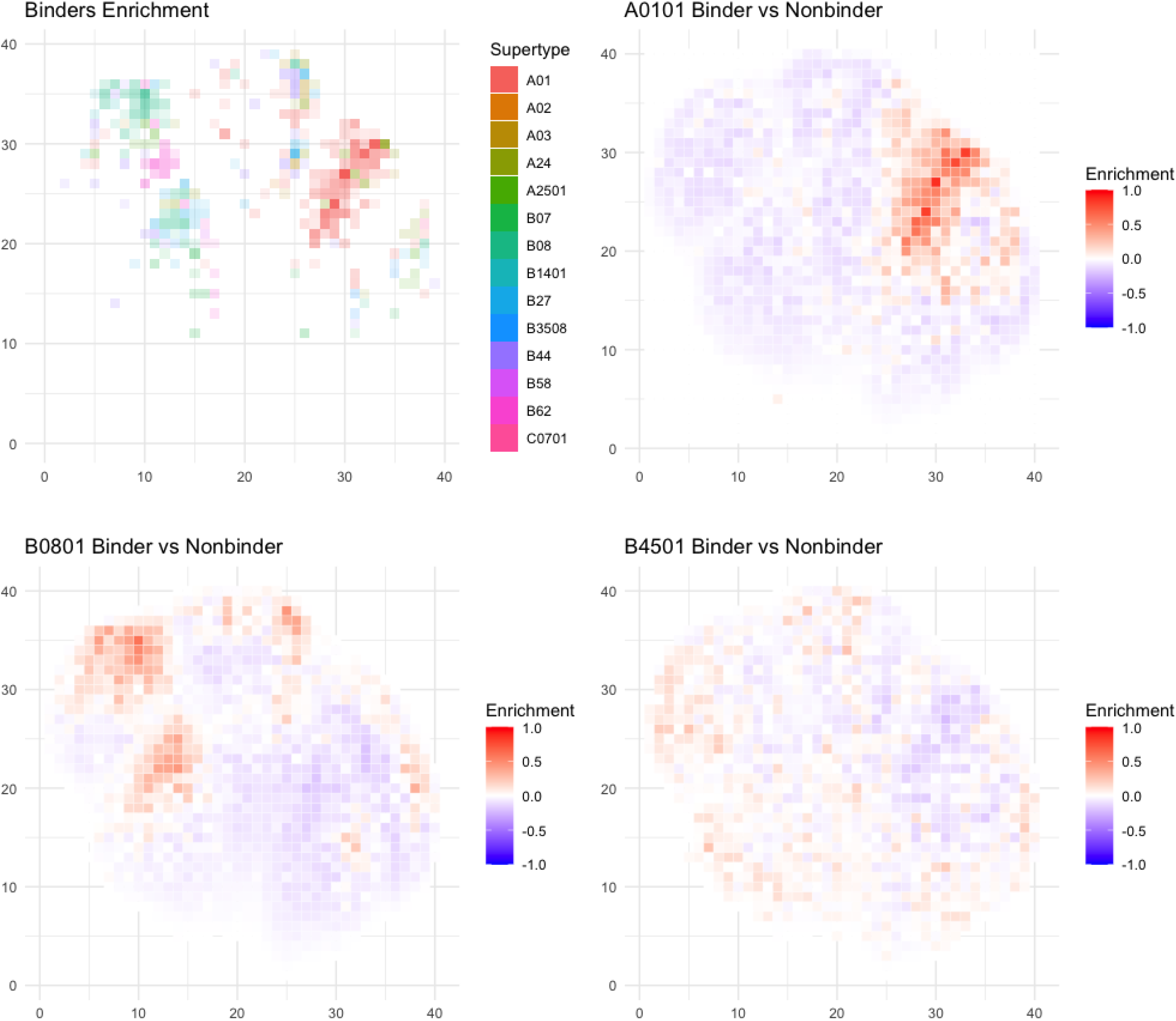
Differential distributions of physical properties for 9-mer peptides predicted to bind to HLA alleles. A) The plotting coordinates represent the first two dimensions of a UMAP transform of peptide physical properties, which is divided into 1600 (40×40) equivalently-sized square bins (see Methods). For each bin where there is at least one HLA allele with >0.2% difference in proportion of all peptides predicted to bind v. non-binders, the identity of the most enriched allele is shaded in the color corresponding to that allele’s supertype as corresponding to the legend. B-D) Example plots of three different alleles (A01:01, B08:01, and B45:01) with different distributions of binders. Each box represents enrichment as the percent peptide difference between predicted binders and non-binders for the given allele. The color scale shows the percent of peptides difference in the given box, with red meaning a larger number of predicted binders and blue meaning a larger number of predicted non-binders.

## DISCUSSION

To the best of our knowledge, this is the first study to examine the consistency of predictions of peptide-MHC binding across different tools, and to explore the quality and quantity of training data in this context. We note several limitations to this work. Firstly, we confined attention to MHC class I peptides and did not include predictions for MHC class II (36), of which there are numerous alleles. We also excluded from consideration any potential contributions of proteasomal cleavage or other antigen processing machinery to MHC binding (37–39). We did not seek to comprehensively assess all available tools for peptide-MHC binding affinity prediction, but rather confined our attention to four of the most widely used tools. The majority of our randomly generated peptides are not known to be found in nature and may not represent the optimal background distribution for measuring allele promiscuity or interrater reliability between tools primarily used for human and pathogenic peptides. While our analysis of peptides leveraged four essential and well-described amino acid physical properties, there may exist unassessed latent features that could capture additional variance and improve dimensionally-reduced comparisons. We did not assess the extent to which mass spectrometry biases in the training datasets might affect peptide-MHC predictions (40–43). Lastly, we did not evaluate individual tool performance based on known epitopes as this has been previously reported (23–27,44–48).

Our work raises fundamental questions about the fidelity of peptide-MHC binding prediction tools. Why, for instance, can predictions be so discordant among tools for which training datasets are otherwise so similar? We especially worry about the real-world use of these prediction tools for alleles without any direct basis in training data. Why is the predicted range of allele promiscuity so substantial, and yet not demonstrative of any meaningful differences in enrichment between potential foreign versus self antigens? Moreover, is this differential promiscuity a universal biological phenomenon, with certain alleles being generally poor functional presenters of antigen? If this is the case, what selective advantage might have evolutionarily maintained these alleles in the population? Evaluating more viruses – as well as bacteria, fungi, and other pathogens – and linking these analyses with metrics such as evolutionary distance may give greater insight into the relationship between HLA evolution and disease.

## METHODS

### Sequence retrieval, peptide filtering, and kmerization

FASTA-formatted protein sequence data was retrieved from the National Center of Biotechnology Information (NCBI) (49,50) using RefSeq as of 1-31-22 for BK, SARS-CoV-2, HHV-5, HHV-6, HSV-1, HSV-2, HSV-4, and Human. Protein sequence data was inputted into netchop v3.0 “C-term” model with a cleavage threshold of 0.1 to remove peptides that were not predicted to undergo canonical MHC class I antigen processing via proteasomal cleavage (of the peptide’s C-terminus). The results from netchop v3.0 were then kmerized sequentially into 8- to 12-mers. Code used for kmerization and netchop filtering can be found at: https://github.com/Boeinco/peptide-MHCassess. We additionally generated a set of 1 million random peptides of length 8-12 drawn uniformly at random. Peptide sets had negligible overlap (<1% shared between human vs viral vs random peptides).

### Peptide-MHC class I binding affinity predictions

MHC class I binding affinity predictions were performed for the peptides generated from the kmerization process above using 4 tools: netMHCpan v4.1 (23), HLAthena v1.0 (27), MHCflurry v2.0 (25), and MHCnuggets v2.3 (51). netMHCpan was run with default options with the ‘-l’ option to specify peptides of lengths 8-12. MHCflurry was run with default options. MHCnuggets was run with default options. HLAthena was run using the dockerized version of HLAthena with default options, which predicts peptides of length 8-11. MHC class I binding affinity predictions were performed for each of 24, 26, and 2, HLA-A, -B, and -C alleles, respectively. Only alleles that were in common between all 4 tools were used (52 total alleles in common between 2489 possible alleles). Binding affinity values were converted to binding probability values for MHCflurry and MHCnuggets using 1-log(binding affinity) / log(50000) in order to match HLAthena and netMHCpan binding probability predictions. Alleles were grouped into supertypes when applicable using the HLA class I revised classification (29).

### Dimensional reduction and binning analysis

Peptides were converted into physical property matrices using amino acid sequence mapping into a 4*kmer length matrix containing each amino acid’s properties in sequence. The following physical properties of the amino acids were encoded: side chain polarity was recorded as its isoelectric point (pI) (52), the molecular volume of each side chain was recorded as its partial molar volume at 37°C (53), the hydrophobicity of each side chain was characterized by its simulated contact angle with nanodroplets of water (54) and conformational entropy was derived from peptide bond angular observations among protein sequences without observed secondary structure (55).

Each dimensional reduction was performed on the pooled set of k-mers. UMAP dimensionality was performed using uwot UMAP R implementation v0.1.11. PCA was performed using default `prcomp()` functions in base R v4.1.3.

For each peptide source, binned matrices were computed using the `bin2()` function with 40×40 (1600) bins from the `Ash` v1.0.15 package (56) in R v4.1.3. Bin values were then divided by the total number of peptides to create bins with the % of total peptides. In order to compare between 2 peptide sources, a matrix, called the difference matrix, is created by subtracting one matrix of a peptide source from another. Taking the absolute value of each bin in the difference matrix, then summing the values together, results in a single metric ranging from 0-2 measuring the difference in binned density between 2 peptide sources, the value 2 indicating that no peptides were shared between bins and the value 0 indicating the same percentage of peptides in every bin (Methods Figure 1).

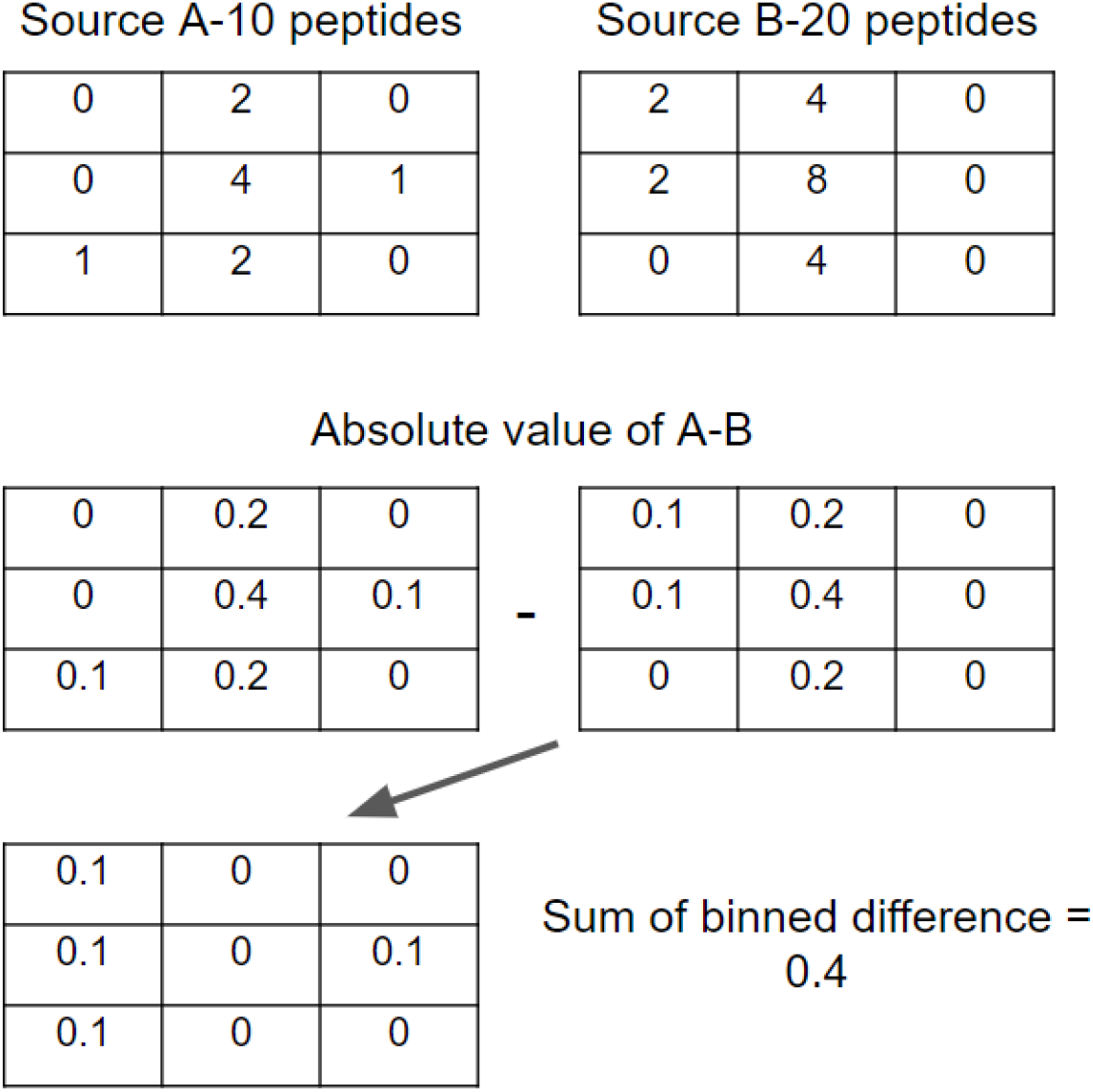
Methods Figure 1.

### Allele ordering similarity

For each allele-peptide source combination, the percentage of peptides predicted to bind with a binding probability score of 0.5 or greater was calculated for all processed peptides. 0.5 binding score is estimated to be equivalent to 250-300nM depending on the tool used. For each peptide source, alleles were ranked from best to worst binders (most to least peptides >= 0.5 score) t. In order to compute allele ordering similarity between 2 peptide sources for a single tool, Spearman’s Rank Correlation Coefficient was calculated between the 2 sets of allele ranks.

For the random group 1 vs random group 2 analysis, we conducted 100 replicates of dividing the randomly generated peptides into 2 random groups and performed a Spearman rank test of allele ordering between these groups for each of the tools.

### Interrater reliability

Intraclass correlation coefficients (ICCs) were calculated using the `ICC()` function from the `IRR` v0.84.1 R package (57). Binding prediction scores for all 1 million randomly generated peptides were separated by tool and HLA allele, and an ICC was calculated as the interrater reliability metric between the 4 tools for each allele. ICC was also between the 4 tools on a per peptide basis, each peptide receiving a score across 4 tools using predictions separated by tool and peptide.

## SUPPLEMENTARY MATERIAL

**Sup Figure 1:**
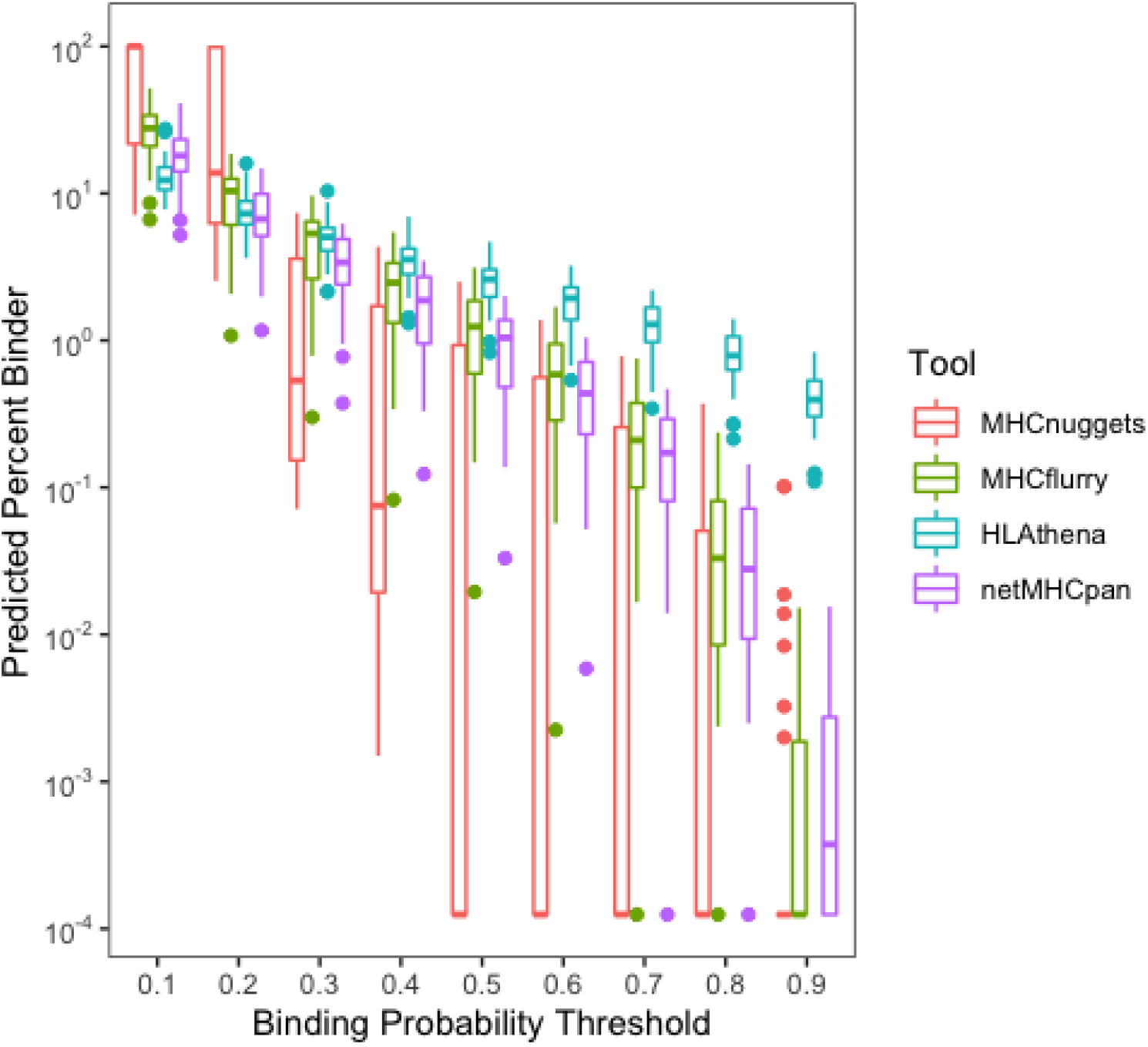
Boxplots of the relationship between predicted binding and the threshold used to determine binding for random peptides. Each color represents a different tool with each boxplot representing the IQR of predicted percent peptides to bind for the given threshold.

**Sup Figure 2.**
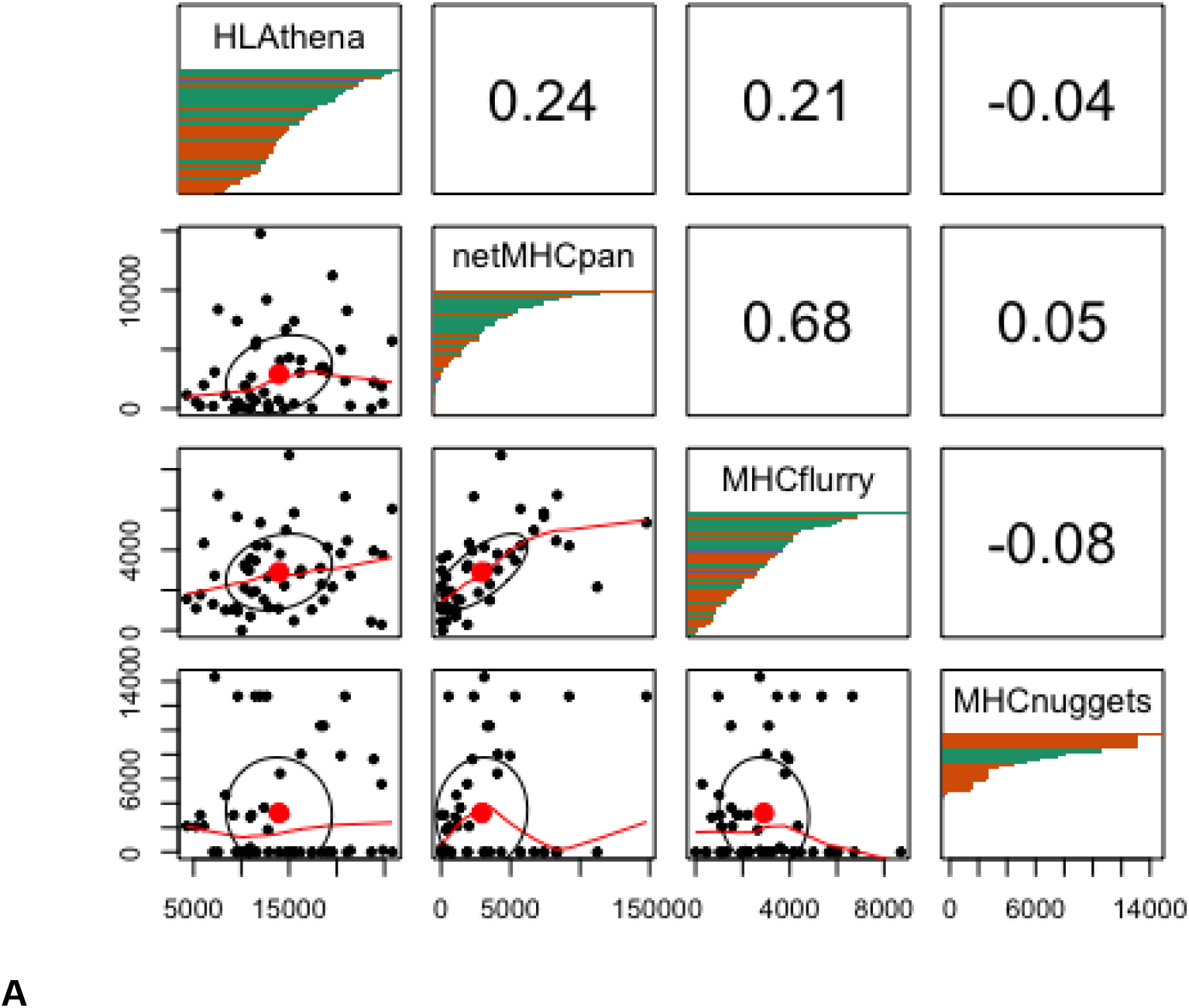
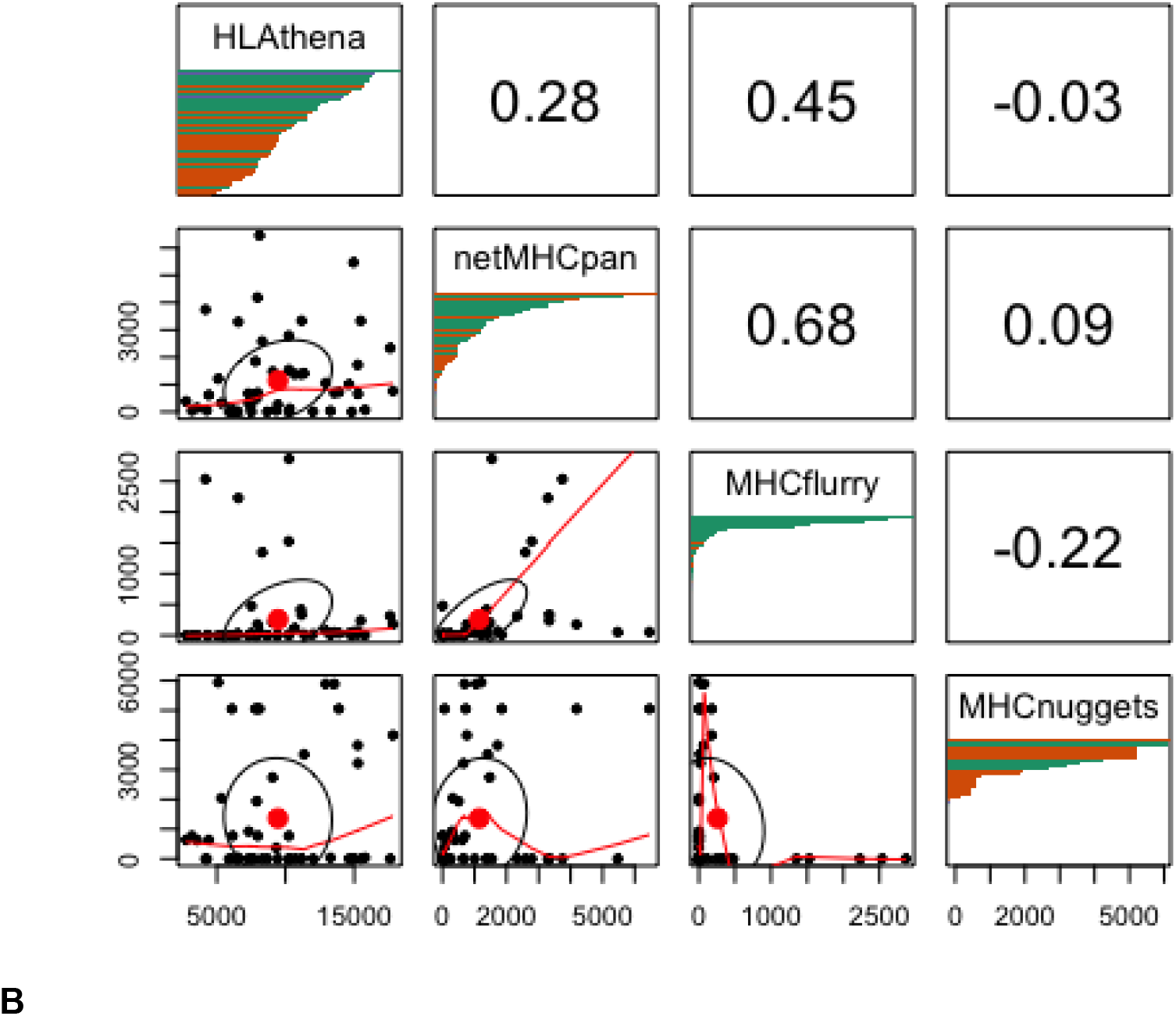
Pairplot of HLA allelic presentation of 8-11mers from the random proteome. The lower left triangle displays scatter plots of peptides predicted to bind using 0.6 (A) and 0.7 (B) as cutoffs respectively between 2 tools with each point representing an HLA allele. The upper right triangle represents the Spearman correlation of the number of peptides predicted to bind to all alleles between tools. Note that MHCnuggets has a number of alleles with 0 random peptides predicted to bind. The diagonal panels show distribution of HLA allelic presentation from the random proteome for each tool. The number of peptides that putatively bind to each of the HLA alleles is shown along the x-axis as a series of horizontal bars with green, orange, and purple colors representing HLA-A, -B, and -C alleles, respectively, sorted in order of decreasing quantity of binders.

**Sup Figure 3:**
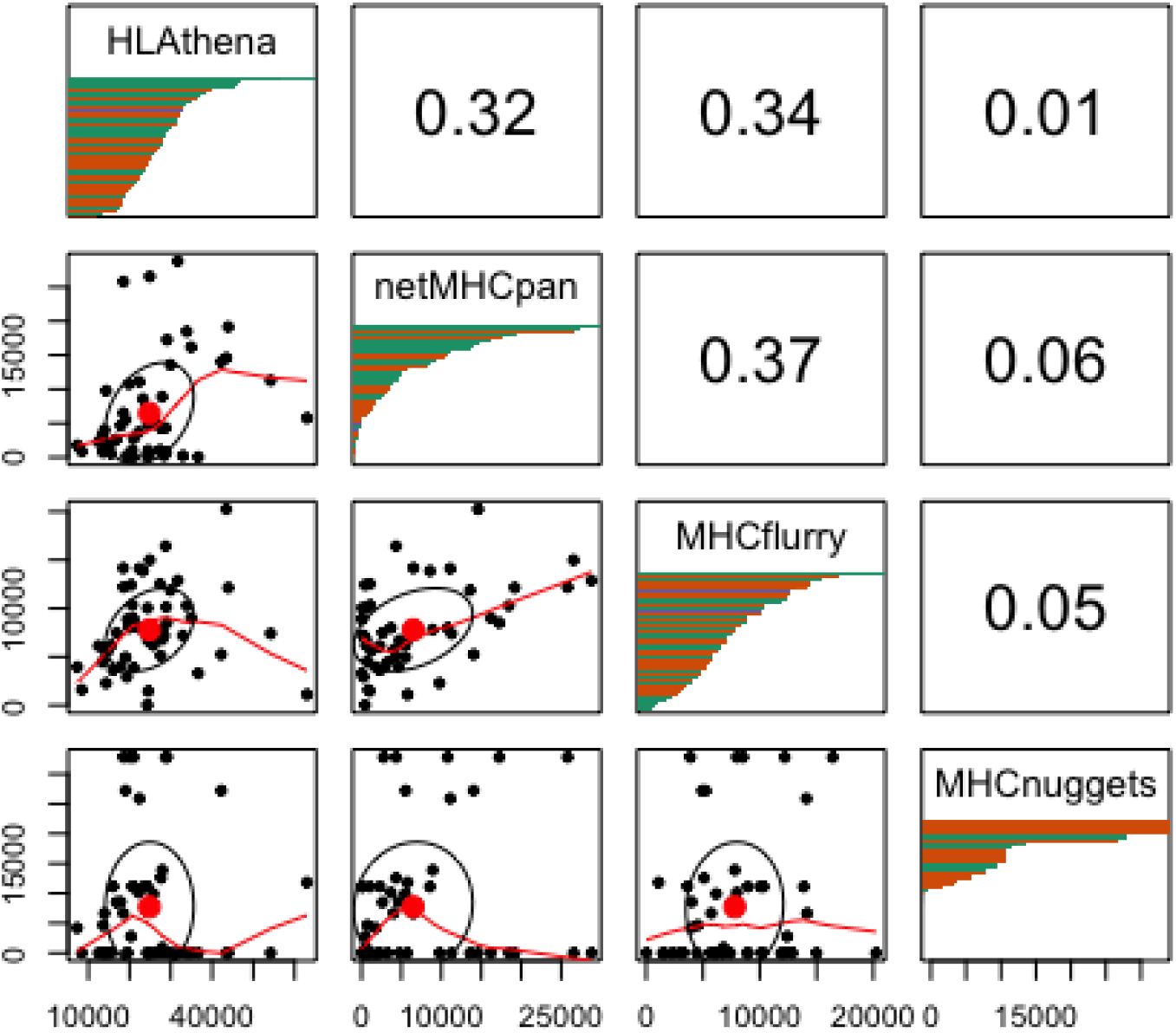
Pairplot of HLA allelic presentation of 8-11mers from the human and viral proteome. The lower left triangle displays scatter plots of peptides predicted to bind (>= 0.5 binding probability score) between 2 tools with each point representing an HLA allele. The upper right triangle represents the Spearman correlation of the number of peptides predicted to bind to all alleles between tools. Note that MHCnuggets has a number of alleles with 0 random peptides predicted to bind. The diagonal panels show distribution of HLA allelic presentation from the random proteome for each tool. The number of peptides that putatively bind to each of the HLA alleles is shown along the x-axis as a series of horizontal bars with green, orange, and purple colors representing HLA-A, -B, and -C alleles, respectively, sorted in order of decreasing quantity of binders.

**Sup Figure 4.**
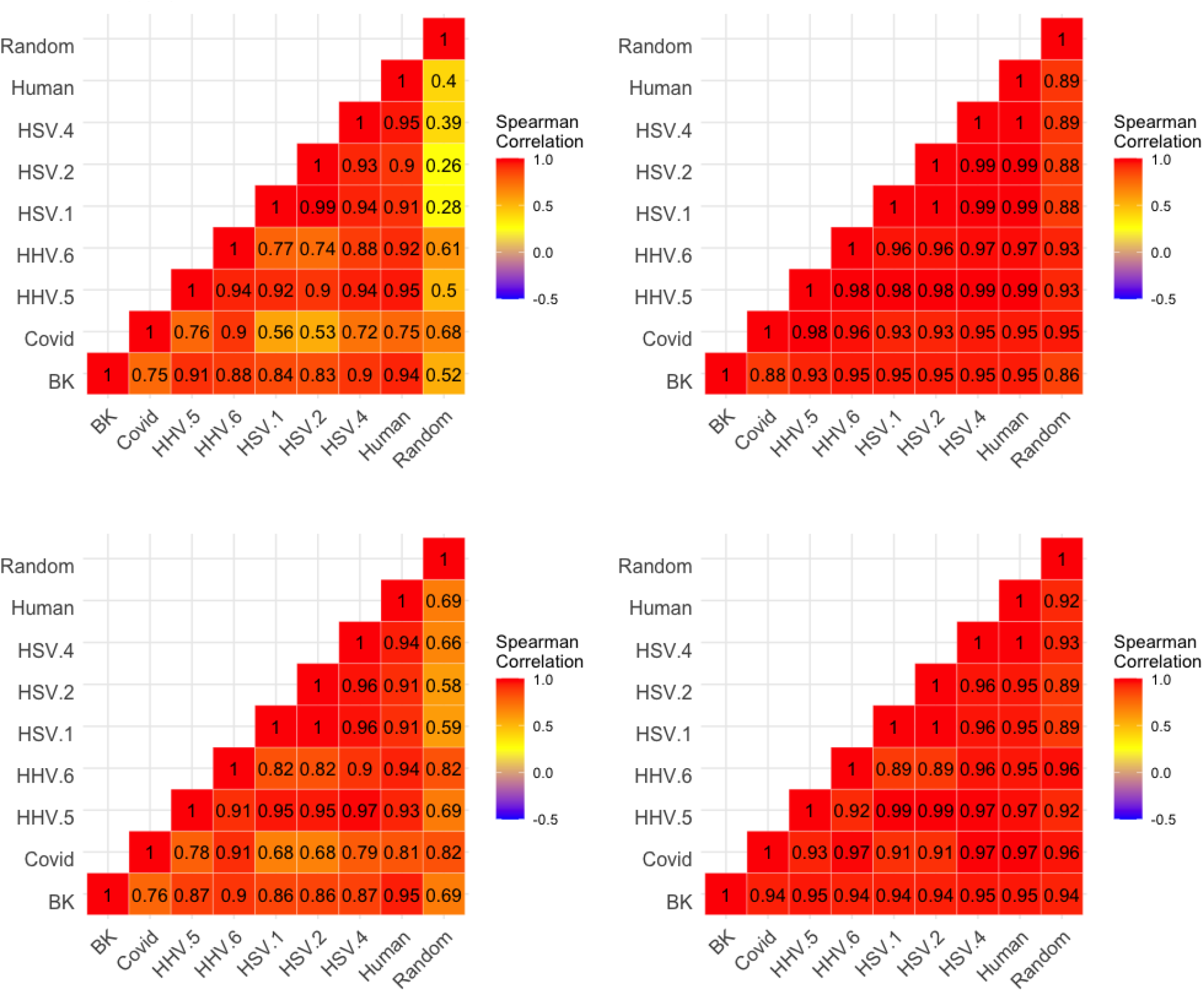
Heatmaps of correlation between peptides for each species of predicted allelic promiscuity across alleles. A) Spearman correlation is shown between peptide sources for HLAthena-based predictions. Analogous data is shown for netMHCpan, MHCflurry, and MHCnuggets in panels B, C, and D, respectively.

**Sup Figure 5.**
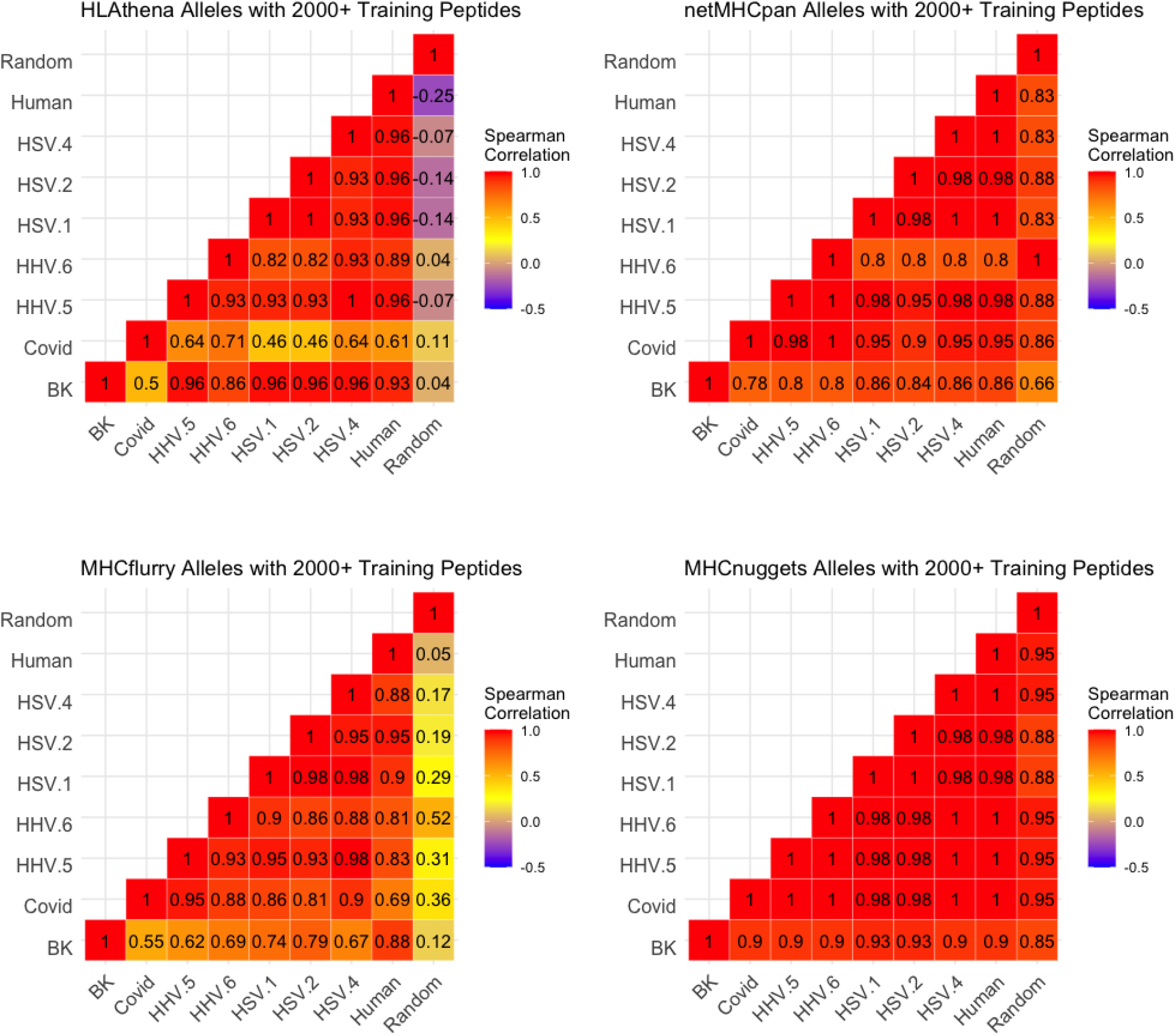
Heatmaps of correlation between peptides for each species of predicted allelic promiscuity across alleles for which there was a minimum of 2000 peptides of training data. A) Spearman correlation is shown between peptide sources for HLAthena-based predictions. Analogous data is shown for netMHCpan, MHCflurry, and MHCnuggets in panels B, C, and D, respectively.

**Sup Figure 6.**
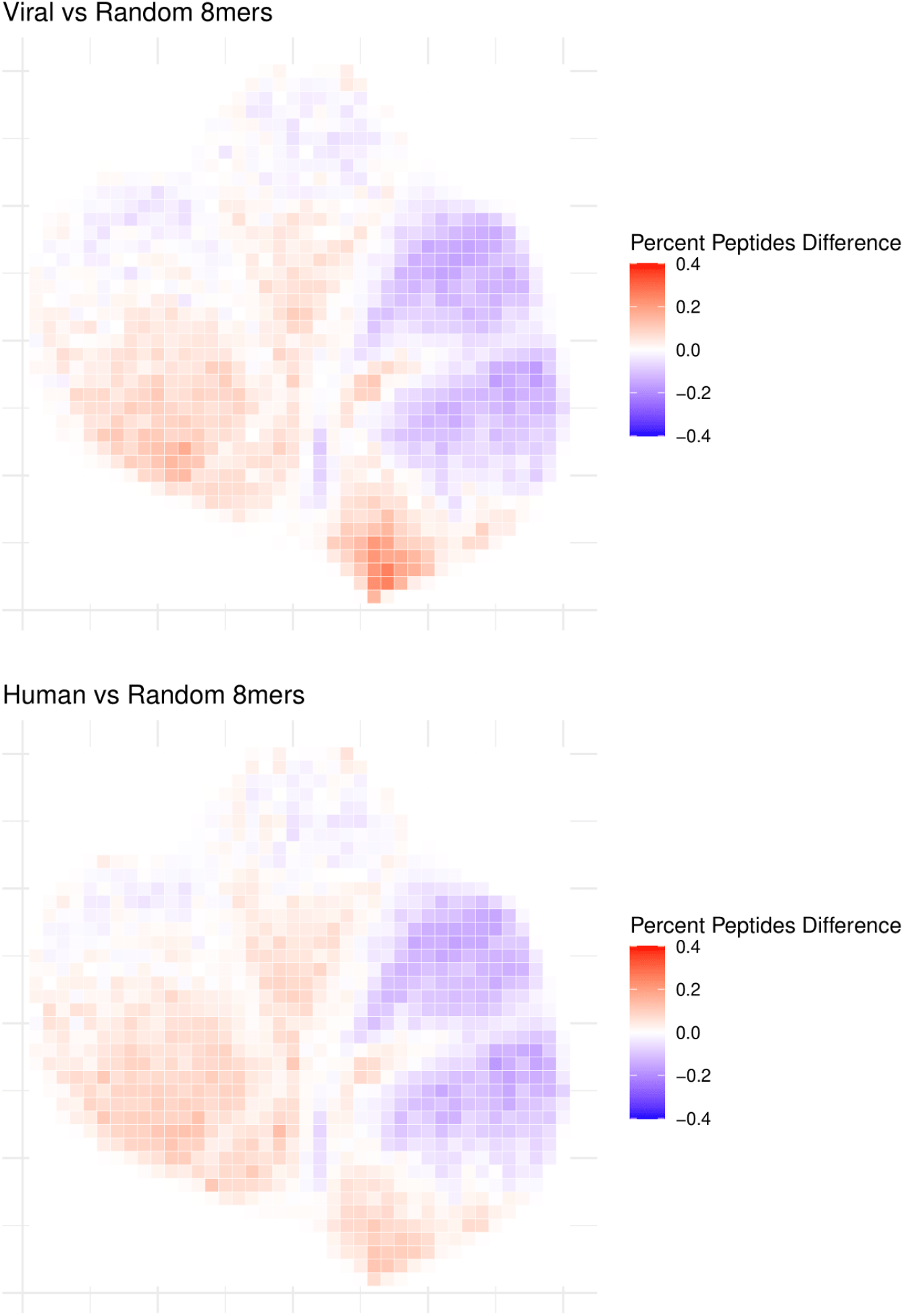
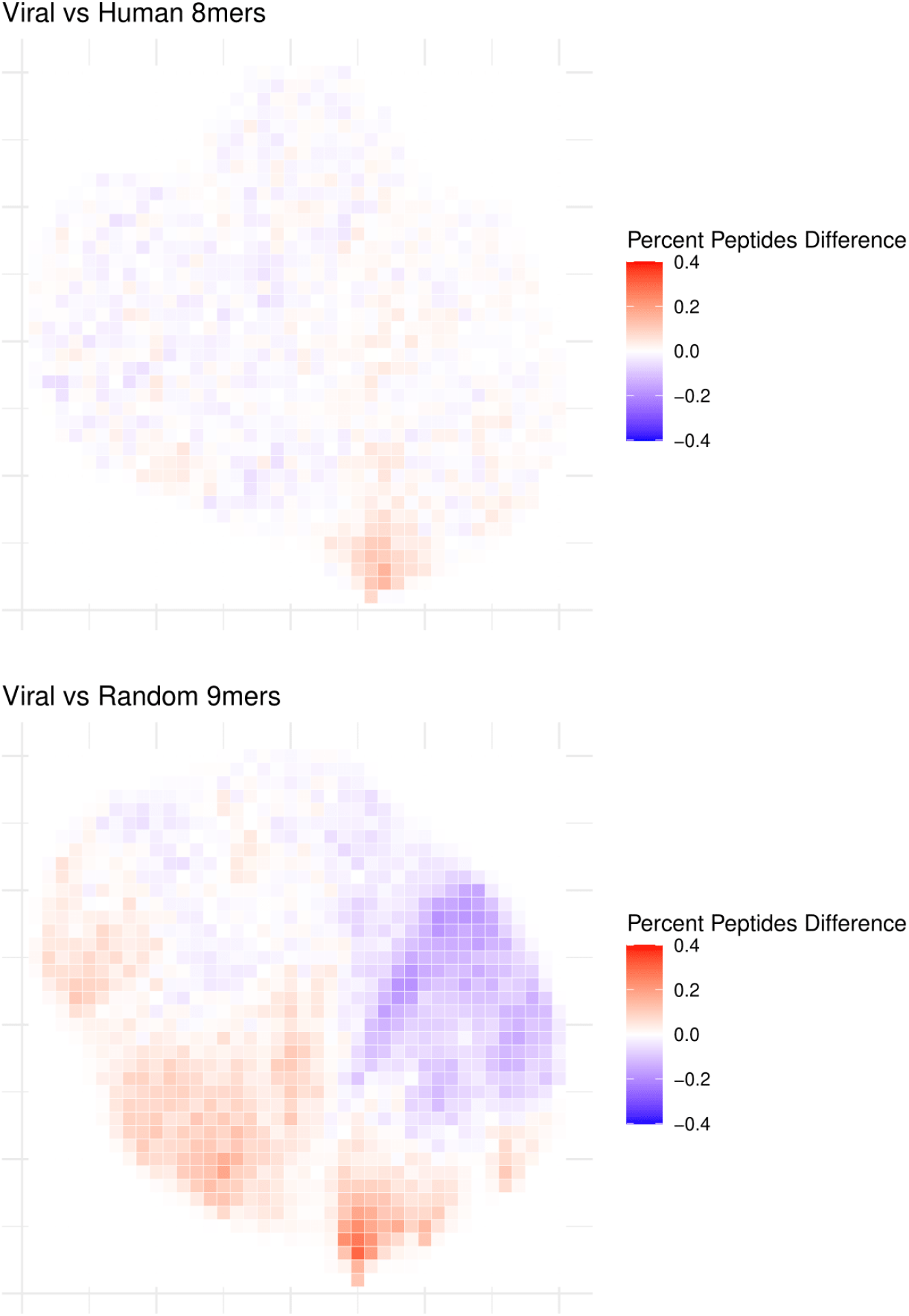
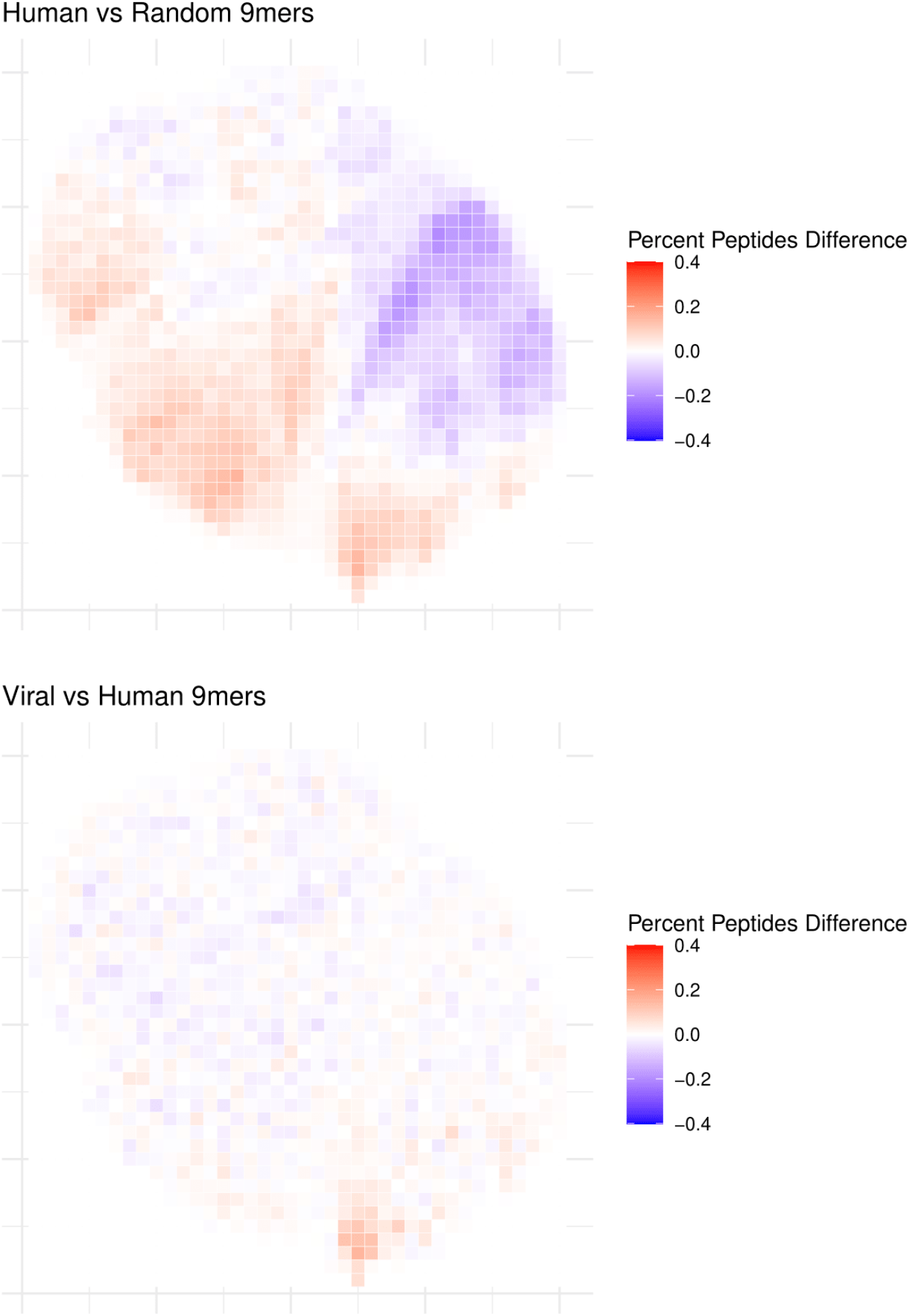
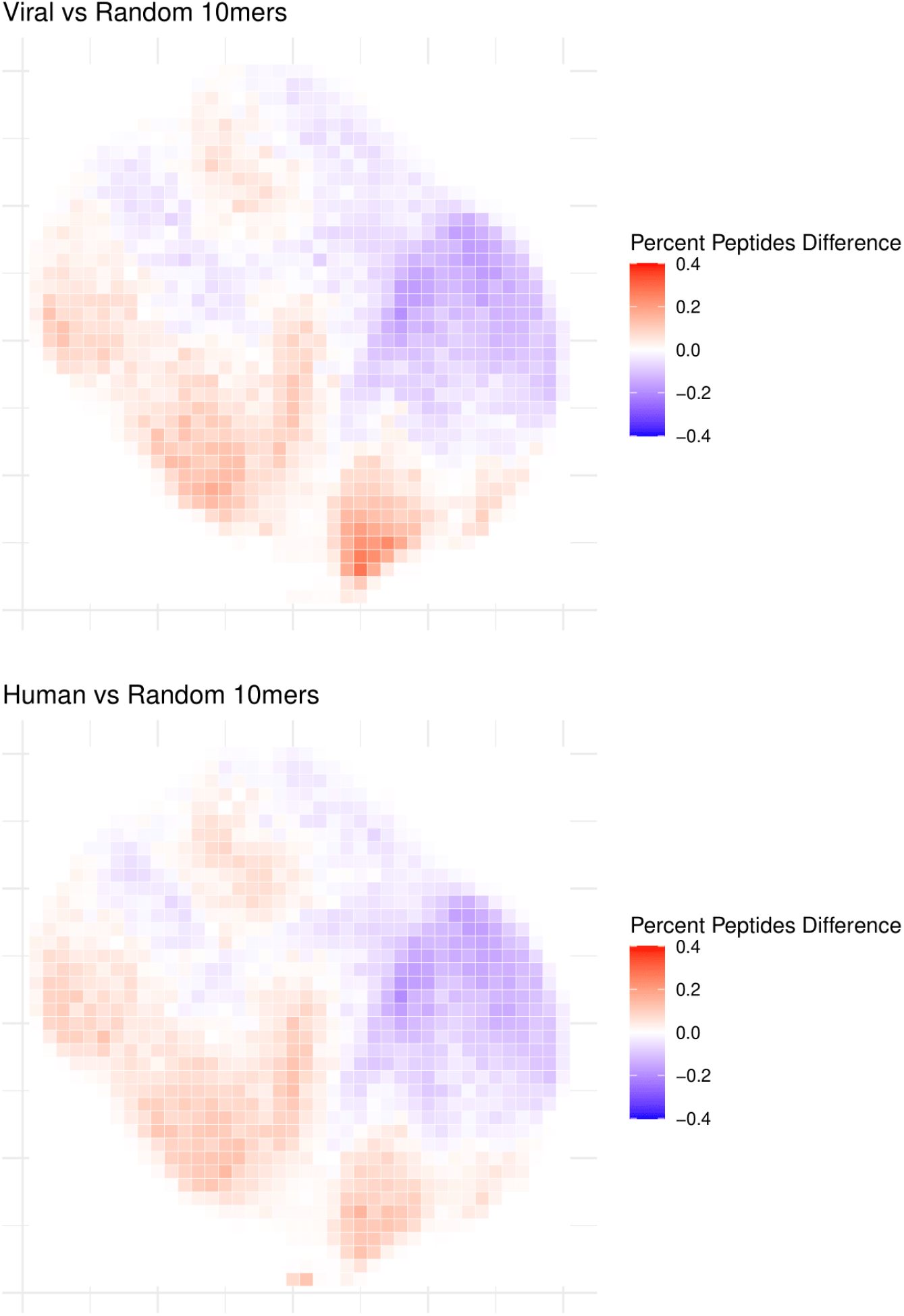
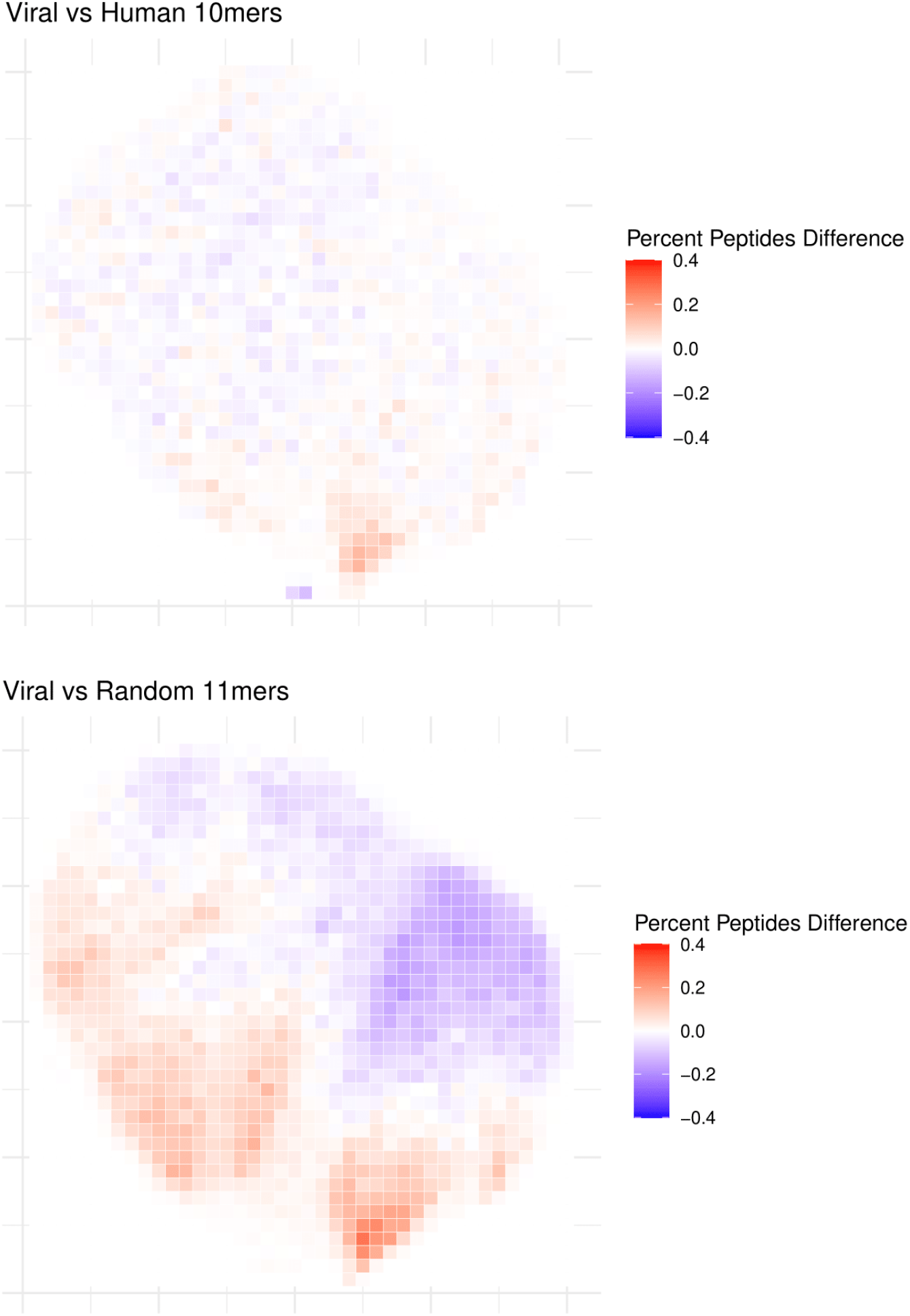
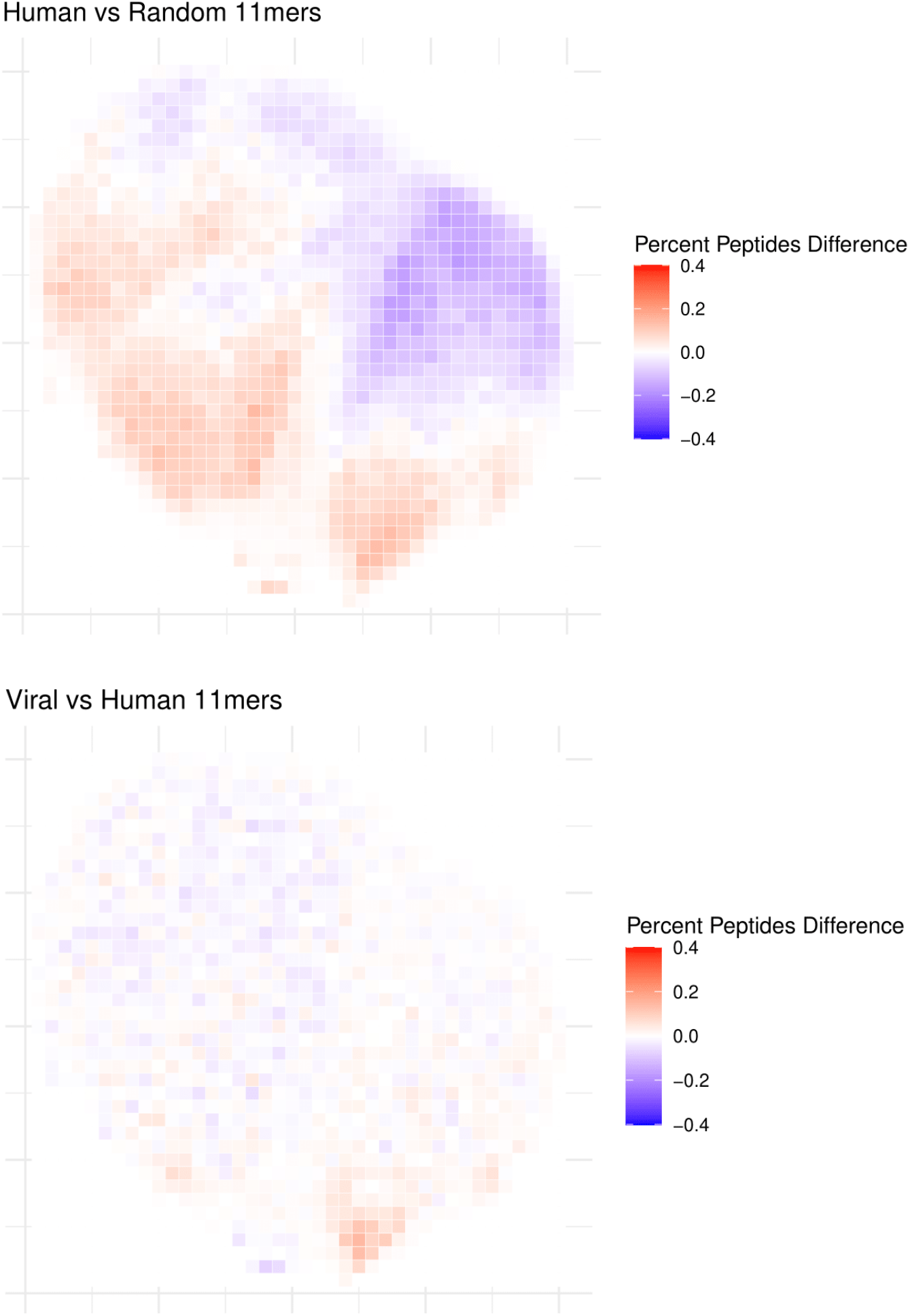
Peptide physical property differences between different peptide sources. Each tile plot is composed of 1600 tiles, with each tile colored by the percent peptide difference between the 2 peptide sources in that particular tile. Red indicates an enrichment of the first label (e.g. viral vs human, viral enrichment will be red) while blue indicates enrichment of the second label.

**Sup Figure 7.**
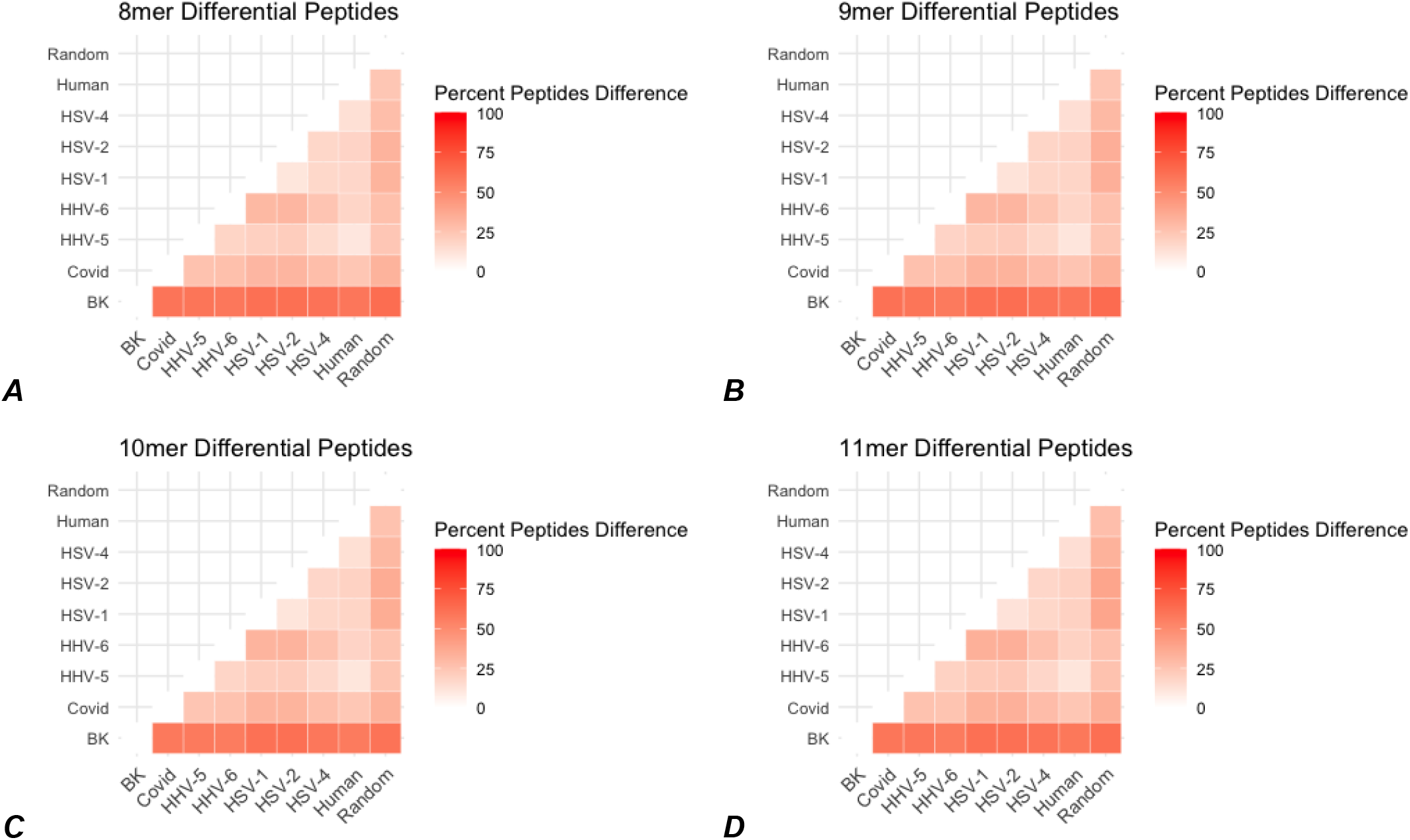
Peptide physical property difference by k-mer length. Each heatmap is the pairwise percent difference metric between each pair of peptide sets. The redder the value, the more difference in the percent difference metric.

**Sup Figure 8.**
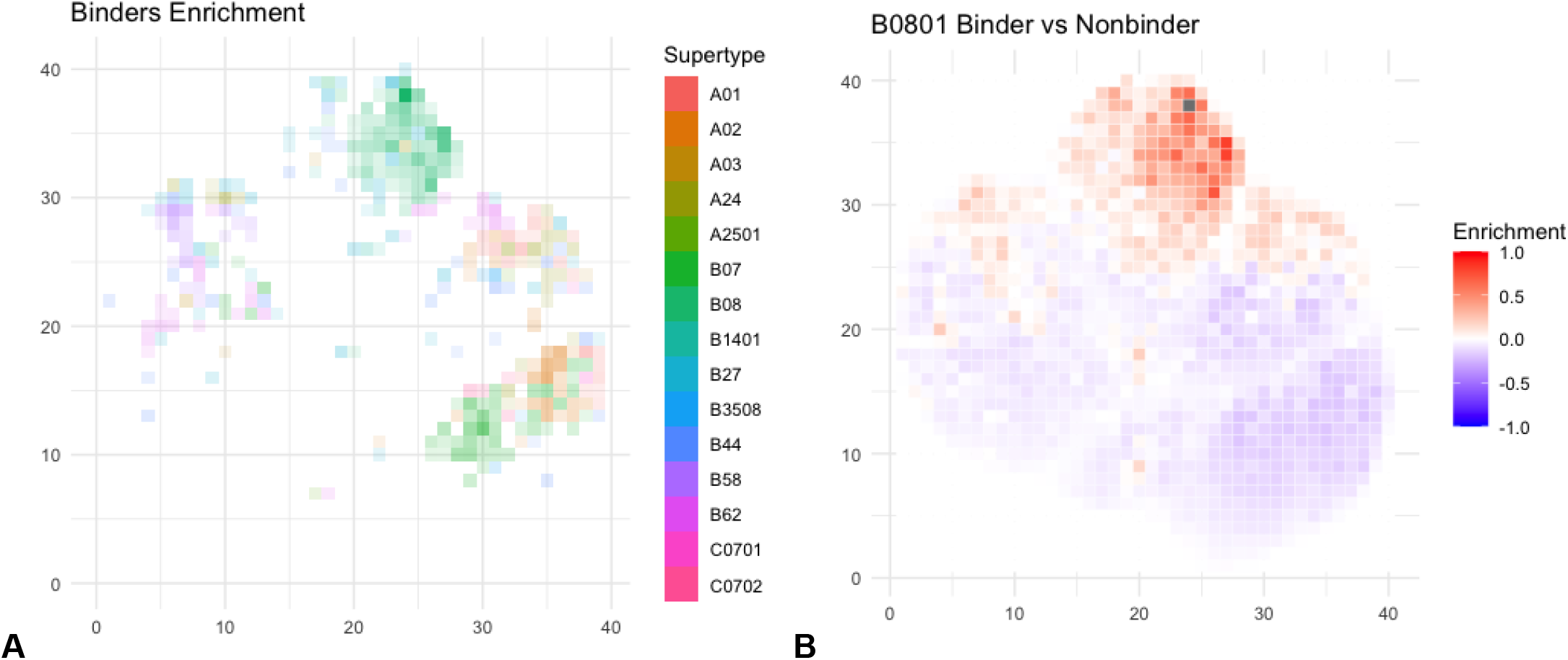
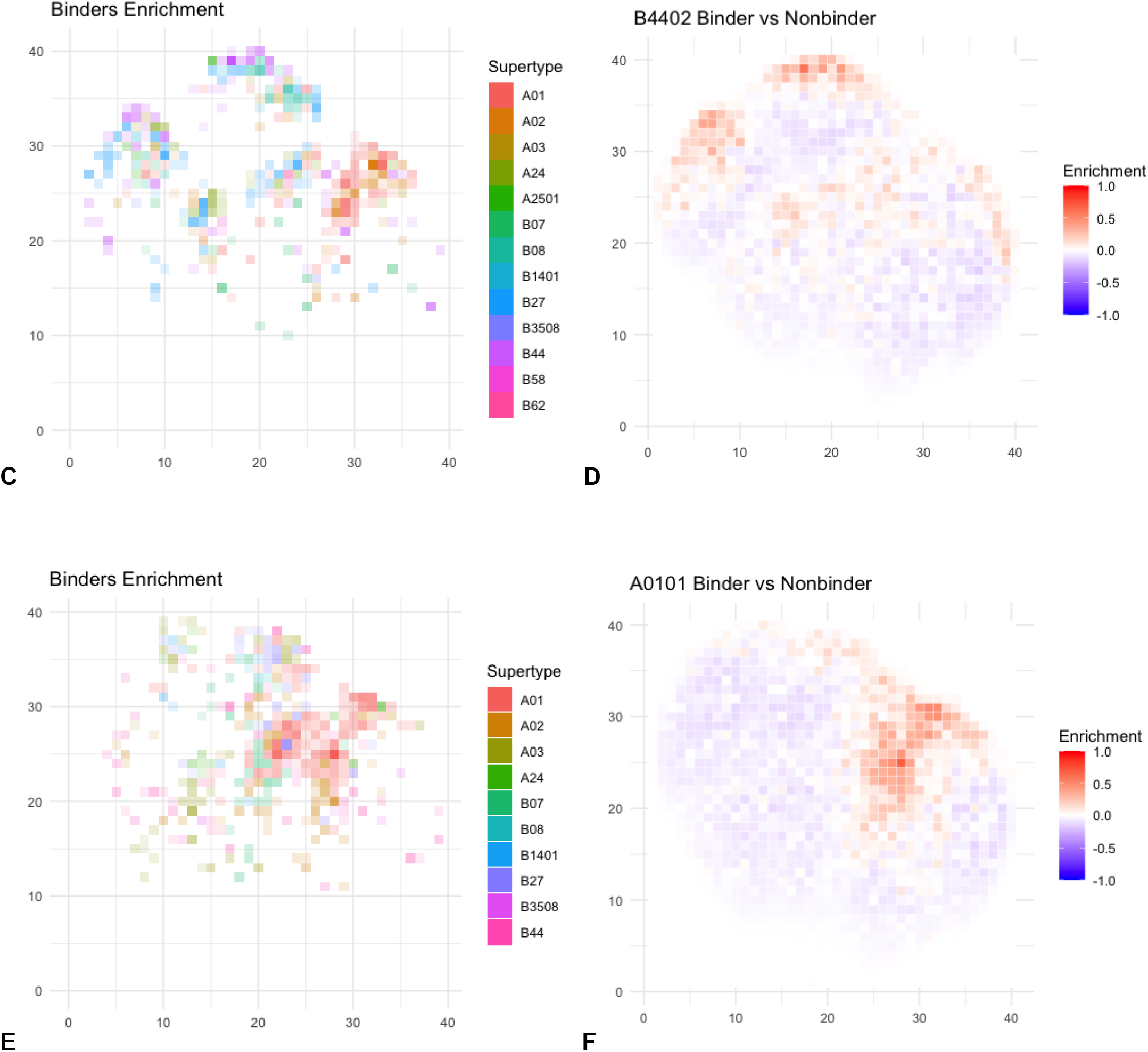
Differential distributions of physical properties for 8,10, and 11-mer peptides predicted to bind to HLA alleles. A,C,E) Tile plots highlighting binders enrichment 8, 10, and 11-mers respectively. The plotting coordinates represent the first two dimensions of a UMAP transform of peptide physical properties, which is divided into 1600 (40×40) equivalently-sized square bins (see Methods). For each bin where there is at least one HLA allele with >0.2% difference in proportion of all peptides predicted to bind v. non-binders, the identity of the most enriched allele is shaded in the color corresponding to that allele’s supertype as corresponding to the legend. B,D,F) Example plots of alleles with different distributions of binders for 8, 10, and 11-mers respectively. Each box represents enrichment as the percent peptide difference between predicted binders and non-binders for the given allele. The color scale shows the percent of peptides difference in the given box, with red meaning a larger number of predicted binders and blue meaning a larger number of predicted non-binders.

## Notes

### Competing Interest Statement

The authors have declared no competing interest.

